# Profile hidden Markov model sequence analysis can help remove putative pseudogenes from DNA barcoding and metabarcoding datasets

**DOI:** 10.1101/2021.01.24.427982

**Authors:** T. M. Porter, M. Hajibabaei

## Abstract

**Background:** Pseudogenes are non-functional copies of protein coding genes that typically follow a different molecular evolutionary path as compared to functional genes. The inclusion of pseudogene sequences in DNA barcoding and metabarcoding analysis can lead to misleading results. None of the most widely used bioinformatic pipelines used to process marker gene (metabarcode) high throughput sequencing data specifically accounts for the presence of pseudogenes in protein-coding marker genes. The purpose of this study is to develop a method to screen for obvious pseudogenes in large COI metabarcode datasets. We do this by: 1) describing gene and pseudogene characteristics from a simulated DNA barcode dataset, 2) show the impact of two different pseudogene removal methods on mock metabarcode datasets with simulated pseudogenes, and 3) incorporate a pseudogene filtering step in a bioinformatic pipeline that can be used to process Illumina paired-end COI metabarcode sequences. Open reading frame length and sequence bit scores from hidden Markov model (HMM) profile were used to detect pseudogenes.

**Results:** Our simulations showed that it was more difficult to identify pseudogenes from shorter amplicon sequences such as those typically used in metabarcoding (∼300 bp) compared with full length DNA barcodes that are used in construction of barcode libraries (∼ 650 bp). It was also more difficult to identify pseudogenes in datasets where there is a high percentage of pseudogene sequences. We show that existing bioinformatic pipelines used to process metabarcode sequences already remove some apparent pseudogenes, especially in the rare sequence removal step, but the addition of a pseudogene filtering step can remove more.

**Conclusions:** The combination of open reading frame length and hidden Markov model profile analysis can be used to effectively screen out obvious pseudogenes from large datasets. There is more to learn from COI pseudogenes such as their frequency in DNA barcode and metabarcoding studies, their taxonomic distribution, and evolution. Thus, we encourage the submission of verified COI pseudogenes to public databases to facilitate future studies.

## Introduction

The mitochondrial cytochrome c oxidase subunit 1 gene, COI, is the official animal barcode marker and large reference databases are available to help identify COI metabarcode sequences from soil, water, sediments, or mixed communities such as those collected from traps [1–3]. Crucially, the COI barcode marker is also a protein coding gene. This is in contrast with the ribosomal DNA markers typically used for marker gene studies of prokaryotes or fungi [4–6]. Until recently, the methodology and bioinformatic pipelines for processing protein coding markers such as COI for animals, the maturase K gene (matK), or the ribulose bisphospate carboxylase large chain gene (rbcL) for plants have been treated in very much the same way, even using the same popular pipelines such as those used to process ribosomal RNA genes.

Innovative methods of processing COI sequence data has arisen in recent years. For example, COI marker analysis need not be limited to operational taxonomic units (OTUs), but may also include the use of exact sequence variant (ESV) analysis for improved taxonomic resolution and permit intraspecific phylogeographic analyses [7–10]. Additionally bioinformatic tools to remove pseudogenes and noise from COI datasets have become available [11–13]. There are currently few options, however, to process COI metabarcode reads that specifically handle COI pseudogenes also known as nuclear encoded mitochondrial sequences (nuMTs). COI pseudogenes have been discussed in the literature largely with regards to COI barcoding efforts and only recently have tools appropriate for handling large batches of COI sequences recently become available [14–17].

Pseudogenes are copies of mitochondrial DNA that have been inserted into the nuclear genome [18]. The mechanism for this is uncertain but may involve the incorporation of mtDNA during the repair of chromosomal double strand breaks [19]. Some mitochondrial pseudogenes are ‘dead on arrival’ due to the different genetic code in the nuclear genome [20]. If the pseudogene has only accumulated a few mutations, the sequence may closely resemble that of a functional COI gene with no frameshift or internal stop codons and may be referred to as a cryptic pseudogene [21]. More apparent pseudogenes, on the other hand, may exhibit stark changes in condon usage bias, transition:transversion ratios, GC content, decreased length, and have unexpected phylogenetic placement [18]. Since the primers used for PCR will bind to paralogous regions in pseudogenes, they will amplify nuMTS in addition to or even preferentially to the target mitochondrial sequence [18, 22]. Including unknown pseudogenes in phylogenetic, biodiversity, or population analyses may introduce noise into analyses, leading to overestimates of haplotype or species richness, or may lead to misleading identifications or relationships [14, 16, 23–26].

The methods needed to detect different types of pseudogenes will vary depending on whether or not many changes have accumulated. Cryptic pseudogenes may be identified by examining raw Sanger chromatograms, similar to looking for evidence for heteroplasmy, by looking for double peaks. The whole gene region may be examined looking for the presence of the control region and stop codon. Conserved regions such as in the inner mitochondrial membrane alpha helices can be examined for changes [27]. More obvious pseudogenes may accumulate substitutions equally in non-synonymous and synonymous regions indicating balanced positive and negative selection at sites across the gene copy or relaxed conservation (dN/dS ratios ∼ 1). This is in contrast with a functional COI gene where substitutions tend to occur in non-synonymous sites so as to preserve amino acid composition and protein structure and dN/dS ratios are expected to be < 1. The result of relaxed purifying selection is the accumulation of indels, frameshifts, and/or the introduction of premature stop codons. The objective of this work is to develop methods to remove such apparent pseudogenes from large COI sequence datasets.

## Bioinformatic Methods

We used three approaches in this study: A) We simulated a DNA barcode dataset by compiling a set of annotated COI genes and pseudogenes from the Barcode of Life Data System (BOLD) and the National Center for Biotechnology Information (NCBI) nucleotide (nt) database for the same set of 10 species; B) We created mock COI metabarcode datasets by mining sequences from BOLD and simulating pseudogenes, and C) We tested a pseudogene filtering method on a previously published freshwater benthos COI metabarcode dataset (Figure 1).

**Figure 1.**
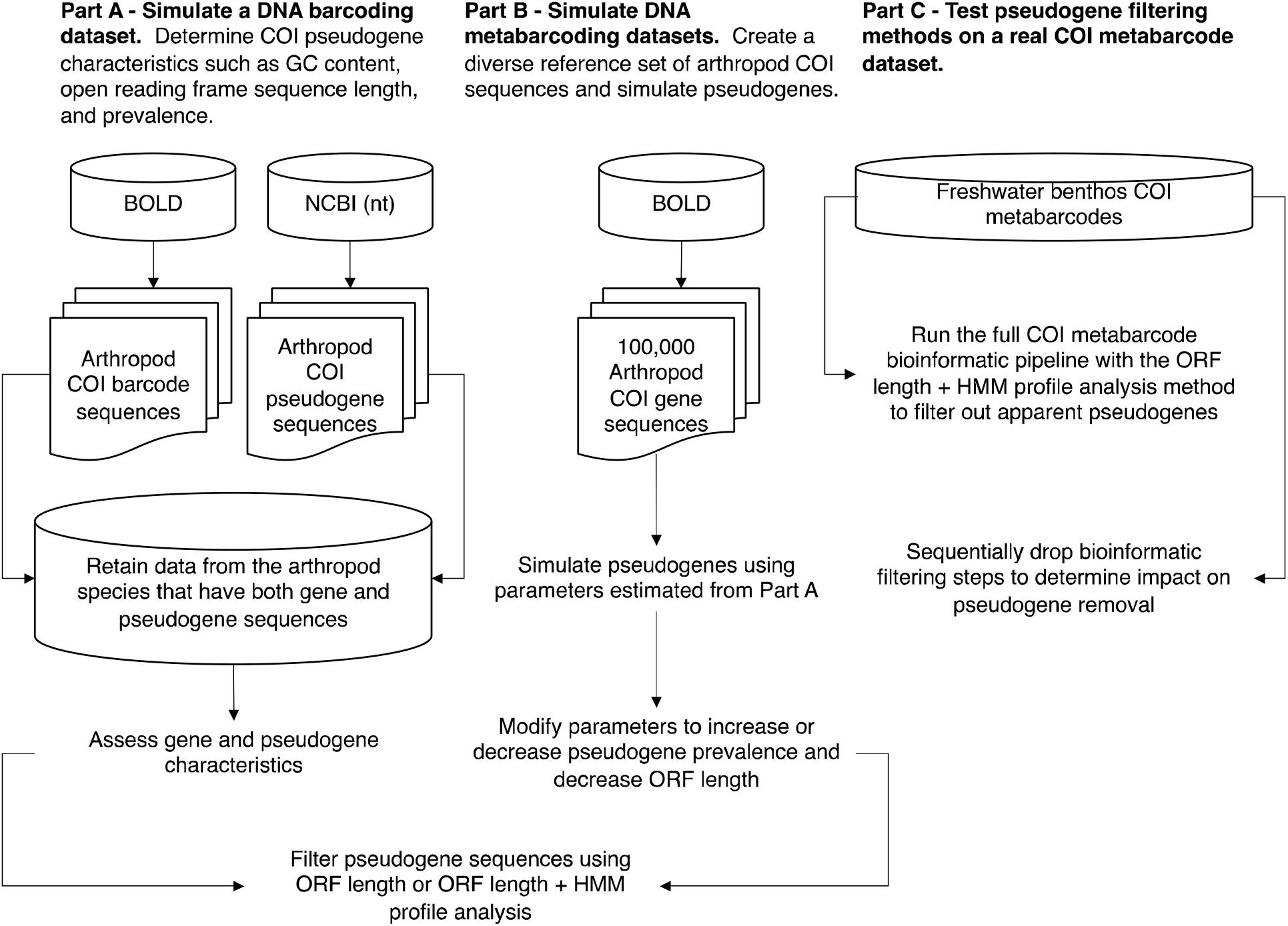
Overview of methods to determine COI pseudogene characteristics and test methods for pseudogene removal. Dataflow for our A) simulated DNA barcode dataset, B) simulated metabarcode datasets, and C) real freshwater COI metabarcode dataset. Abbreviations: BOLD = Barcode of Life Data System; COI = cytochrome c oxidase subunit I mtDNA gene; HMM = hidden Markov model; NCBI = National Centre for Biotechnology Information; nt = nucleotide; ORF = open reading frame.

### Part A: Simulating a DNA barcoding dataset

To simulate a DNA barcoding dataset where multiple sequences are generated for the same species, we retrieved high quality sequences from BOLD and known pseudogenes mined from the NCBI nucleotide database for the same set of species. Sequences from the BOLD data releases were obtained from http://v3.boldsystems.org/index.php/datarelease. Nucleotide sequences for arthropods were selected, ensuring that there were no ambiguities in the nucleotide sequences. If either the nucleotide sequence or amino acid sequence were missing, then the record was discarded. A FASTA file containing arthropod COI pseudogenes was obtained from the NCBI nucleotide database using an Ebot script with the search term “Arthropoda[ORGN] AND pseudogene[TITL] AND (COI[GENE] OR CO1[GENE] OR coxI[GENE] OR cox1[GENE]) AND 50:2000[SLEN]”.[28] A few records had to be edited by hand to isolate the sequence region associated with the COI pseudogene. We retrieved 481 COI nucleotide sequences from BOLD and 112 COI pseudogene nucleotide sequences from the NCBI nucleotide database from the same 10 species (Table 1). This dataset is further described in Table S1 showing proportion of pseudogenes, average length, and average GC content. On average, the length and GC content of pseudogenes from these 10 species are slightly shorter and lower, respectively, than for COI gene sequences.

**Table 1:** Summary of a simulated DNA barcoding dataset containing known arthropod COI pseudogenes

| Class | Order | Species | Gene sequences (% of total) | Pseudogene sequences (% of total) | Subtotals |
| --- | --- | --- | --- | --- | --- |
| Insecta | Coleoptera | *Xylosandrus germanus* | 33 | 1 | 34 |
| Insecta | Hemiptera | *Bemisia tabaci* | 252 | 7 | 259 |
| Insecta | Hemiptera | *Trialeurodes vaporariorum* | 3 | 1 | 4 |
| Insecta | Hemiptera | *Triatoma dimidiata* | 9 | 1 | 10 |
| Insecta | Hymenoptera | *Ectatomma gibbum* | 6 | 1 | 7 |
| Insecta | Hymenoptera | *Halictus rubicundus* | 29 | 2 | 31 |
| Insecta | Hymenoptera | *Melissotarsus insularis* | 135 | 79 | 214 |
| Insecta | Orthoptera | *Cyphoderris monstrosa* | 7 | 14 | 21 |
| Collembola | Entomobryomorpha | *Lepidocyrtus cyaneus* | 5 | 1 | 6 |
| Malacostraca | Decapoda | *Goneplax rhomboides* | 2 | 5 | 7 |
|  |  | Subtotals | 481 (81) | 112 (19) | 593 |

GC content for COI gene and pseudogene sequences were assessed in R using the ‘seqinr’ package [29]. We pooled all the sequences together, then proceeded to filter out just the pseudogene sequences using two different methods:

The first method we used to remove pseudogenes involved screening out ESVs with outlier open reading frame lengths that were very short or very long (SCVUC v4.1.0). This was done by translating arthropod ESVs using ORFfinder v0.4.3 into every possible open reading frame on the plus strand, ignoring nested ORFs, minimum length set to 30. The longest nucleotide (nt) ORFs were retained. Outliers, putative pseudogenes or genuine sequences with PCR/sequencing errors, were identified as sequences shorter than the 25^th^ percentile ORF length - (1.5 * interquartile length) and longer than the 75^th^ percentile ORF length + (1.5 * interquartile length).

The second method we used to remove pseudogenes involved profile hidden Markov model (HMM) analysis (SCVUC v4.3.0). This was done by creating a profile HMM based on BOLD arthropod barcode sequences using HMMER v3.3 available from http://hmmer.org. From the BOLD data releases iBOL phase 0.50 to 6.50, we retrieved all arthropod barcodes 600-700 bp in length. We sorted these sequences by decreasing length using the ‘sortbylength’ command in VSEARCH. We reduced the dataset size by clustering by 80% sequence similarity using the ‘cluster_size’ command and retaining the centroids sequences. As described above, arthropod ESVs were translated and the longest open reading frames were retained for both nucleotide and amino acid (aa) sequences. The amino acid ORFs were aligned with MAFFT v7.455 using the ‘auto’ setting [30]. The nucleotide ORFs were also mapped to the amino acid alignment using TRANALIGN (EMBOSS v6.6.0.0) specifying the invertebrate mitochondrial genetic code [31]. The FASTA file comprised of 6,162 amino acid sequences was converted to Stockholm format. This reference alignment was turned into a model that describes the probabilities for travelling a path along the length of the alignment that moves through match, insert, or deletion states. HMMER was used to build this nucleotide arthropod COI profile hidden markov model (HMM) using the ‘hmmbuild’ command. The HMM was indexed using the ‘hmmpress’ command. Individual arthropod amino acid ORFs were then compared with the profile HMM using the ‘hmmscan’ command. One of the hmmscan outputs is a log odds ratio score (bit score) that compares the likelihood of the query sequence given the model to the likelihood of the query sequence given a random sequence model. When a COI gene is used as the query, we expected a high bit score; whereas when an obvious COI nuMT is used as the query, we expected a low bit score. In this way, putative pseudogenes or genuine sequences with PCR/sequencing errors were identified as amino acid ORFs with short outlier HMMER scores.

We also calculated the number of substitutions per non-synonymous and synonymous sites. Gene sequences and pseudogene sequences were analyzed separately as follows: Amino acid ORFs were aligned using MAFFT v7.455 using the ‘auto’ setting. A codon alignment was created using TRANALIGN (EMBOSS) by mapping the nucleotide ORFs to the amino acid alignment using the invertebrate mitochondrial genetic code. We used the package ‘ggplot2’ in Rstudio to create all plots [32–34]. We used the ‘seqinr’ function ‘kaks’ to calculate the number of substitutions for non-synonymous and synonymous sites [29]. Before calculating dN/dS ratios, we excluded pairwise sequence comparisons where the number of substitutions per synonymous site was < 0.01 (sequences too similar to yield reliable dN/dS) or > 2 (too many substitutions, near saturation, to yield a reliable dN/dS).

To assess how pseudogene sequences could be (mis)identified using the top BLAST hit method, we used the megablast algorithm to find the most similar sequence in the NCBI nucleotide sequence database [35]. We used this method to verify that the expected species was a top match (skipping over the top match if it was the same as the query sequence or if it was an obvious contaminant) and whether or not the top match was to a gene or pseudogene sequence in the reference database. To further visualize phylogenetic divergence between gene and pseudogene sequences for each species, we aligned nucleotide sequences with MAFFT using the ‘auto’ setting. The ‘fdnadist’ Phylip method in the EMBOSS package was used to calculate distances using the Kimura 2-parameter (K2P) model of nucleotide sequence evolution [36, 37]. A neighbor joining tree was saved in Newick format using the ‘fneighbor’ Phylip method in EMBOSS. Statistical support at nodes was calculated by bootstrapping the multiple sequence alignment 1000 times using the ‘fseqboot’ Phylip method in the EMBOSS package then K2P distances and neighbor joining trees were constructed as described above. A majority rule consensus tree was constructed using the Phylip program ‘consense’ [37]. Bootstrap values from the consensus tree were mapped to the phylogram using TreeGraph2 v2.15.0-887 [38]. The tree was mid-point rooted and nodes rotated or collapsed where necessary to improve readability using FigTree v1.4.4 available from http://tree.bio.ed.ac.uk/software/figtree/. Further minor editing to improve readability was performed using Inkscape v1.0.1 available from https://inkscape.org/.

### Part B: Simulating community sequence data

To test our pseudogene filtering methods on a more taxonomically diverse community of arthropods, we performed a simulation study. We created an arthropod COI community based on 100,000 sequences randomly sampled from BOLD. We manipulated this mock community in different ways described below. In our first mock community, based on our simulated DNA barcoding results from Part A where ∼ 19% of our dataset represented pseudogenes, we decided to introduce mutations into 19% of the BOLD sequences. Also based on the results from Part A, we reduced the GC content in our simulated pseudogenes by 2.5% by replacing G/C bases with an A/T bases. In our second mock community, we inserted or deleted bases to introduce frameshift mutations and premature stop codons. To keep the rate of pseudogenization the same as the first mock community, we introduced indels in 2.5% of the bases in our simulated pseudogenes. In the third mock community, we split COI barcode sequences in half to test whether our pseudogene filtering approach would work on shorter barcode sequences similar in length to those generated in COI metabarcoding studies (∼ 300 bp). In a fourth mock community, we doubled the proportion of pseudogenes in the mock community from 19% to 38%. In the fifth mock community, we halved the proportion of pseudogenes in the mock community from 19% to 9.5%. Each of these datasets is further described in Table S1 showing proportion of pseudogenes in the community, average length, and average GC content.

### Part C: Test pseudogene filtering methods using a real COI metabarcode dataset

We used a previously published freshwater benthos COI metabarcode dataset to test our bioinformatic pipeline and two different pseudogene removal strategies [39]. We chose this dataset because it includes results from six different COI amplicons (BR5 [B, ArR5] ∼ 310 bp, F230R [LCO1490, 230_R] ∼ 229 bp, ml-jg [mlCOIintF, jgHCO2198] ∼ 313 bp, BF1 [BF1, BR2] ∼ 316 bp, BF2 [BF2, BR2] ∼ 421 bp, fwh1 [fwhF1, fwhR1] ∼ 178 bp) currently used in a variety of labs in the freshwater COI metabarcode literature [40–47]. The primers and their target taxa are listed in Table S2. Each amplicon covers sites across the COI barcoding region and the mode length ranges from 178 bp (fwh1) to 421 bp (BF2), averaging ∼ 300 bp. The F230R and fwh1 amplicons align to the 5’ end of the barcoding region and the BR5, ml-jg, BF1, and BF2 amplicons align to the 3’ end of the barcode region.

A COI metabarcoding bioinformatic pipeline, SCVUC v4.3.0, was used to process Illumina paired-end reads to output a set of taxonomically assigned ESVs (available from GitHub at https://github.com/Hajibabaei-Lab/SCVUC_COI_metabarcode_pipeline) (Fig 2). This pipeline runs in a conda environment using a snakemake pipeline. Conda is an environment and package manager [48]. It allows most programs and their dependencies to be installed easily and shared with others. Snakemake is a python-based workflow manager [49]. The snakefile contains the commands need to run a bioinformatic pipeline. The configuration file allows users to adjust parameter settings.

**Fig 2.**
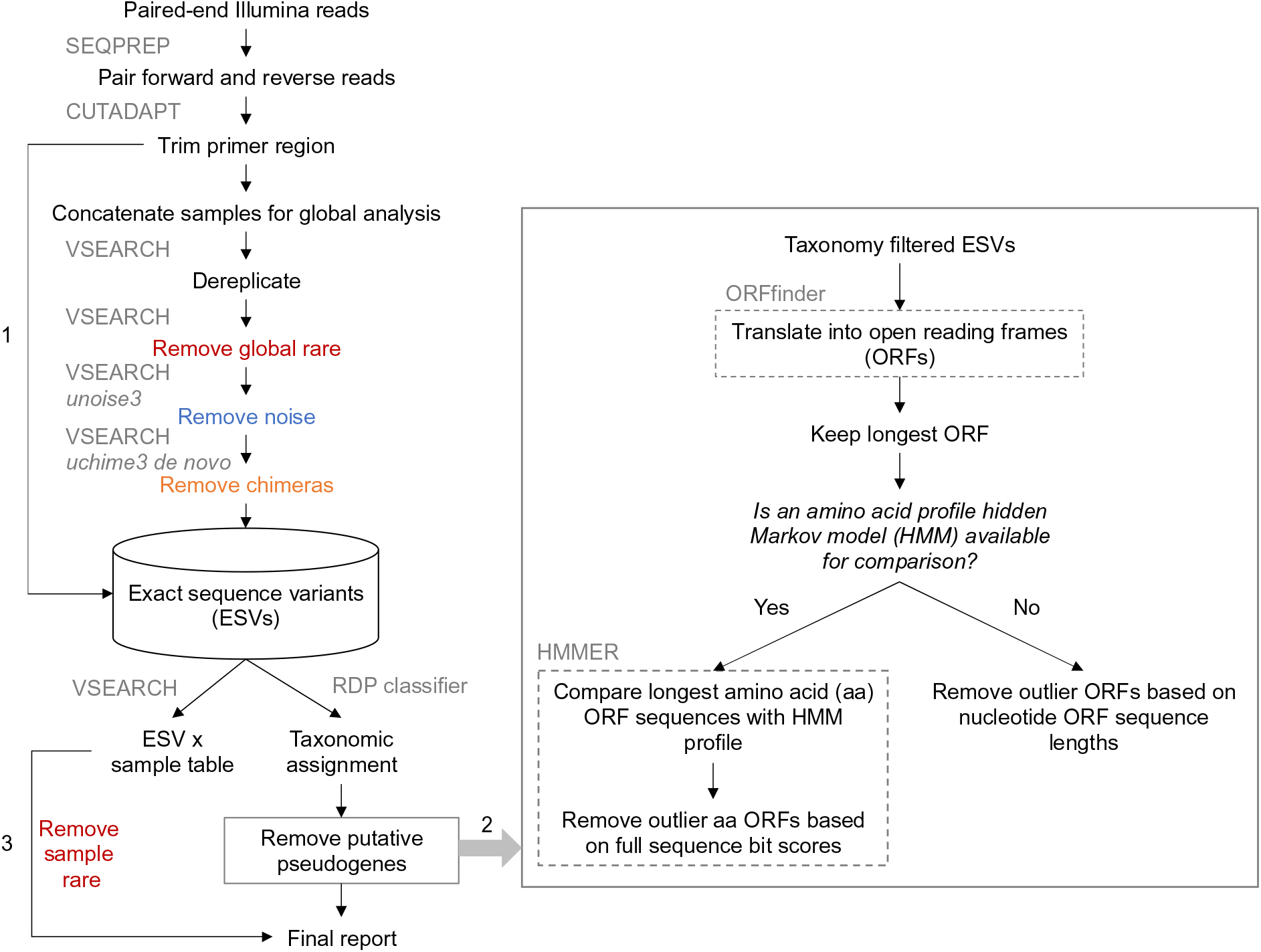
Overview of metabarcoding bioinformatic pipeline that removes apparent pseudogenes. The SCVUC pipeline begins with Illumina paired-end reads. Arrow 1 indicates where globally rare sequence clusters are removed and quality trimmed reads are mapped to denoised exact sequence variants (ESVs) to create a sample x ESV table that contains read numbers. Arrow 2 indicates where pseudogenes can be removed using two different approaches. The first method translates ESVs, retains the longest nucleotide open reading frame (ORF), then removes sequences with very small or very large outlier lengths. The second method translates ESVs, retains the longest amino acid open reading frame, does a profile HMM analysis, then removes sequences with very small outlier full sequence bit scores. Arrow 3 indicates where rare sequence clusters from each sample are removed and read numbers are mapped to the final report. The final report contains all ESVs for each sample, read numbers, ORF sequences, and taxonomic assignments with bootstrap support values.

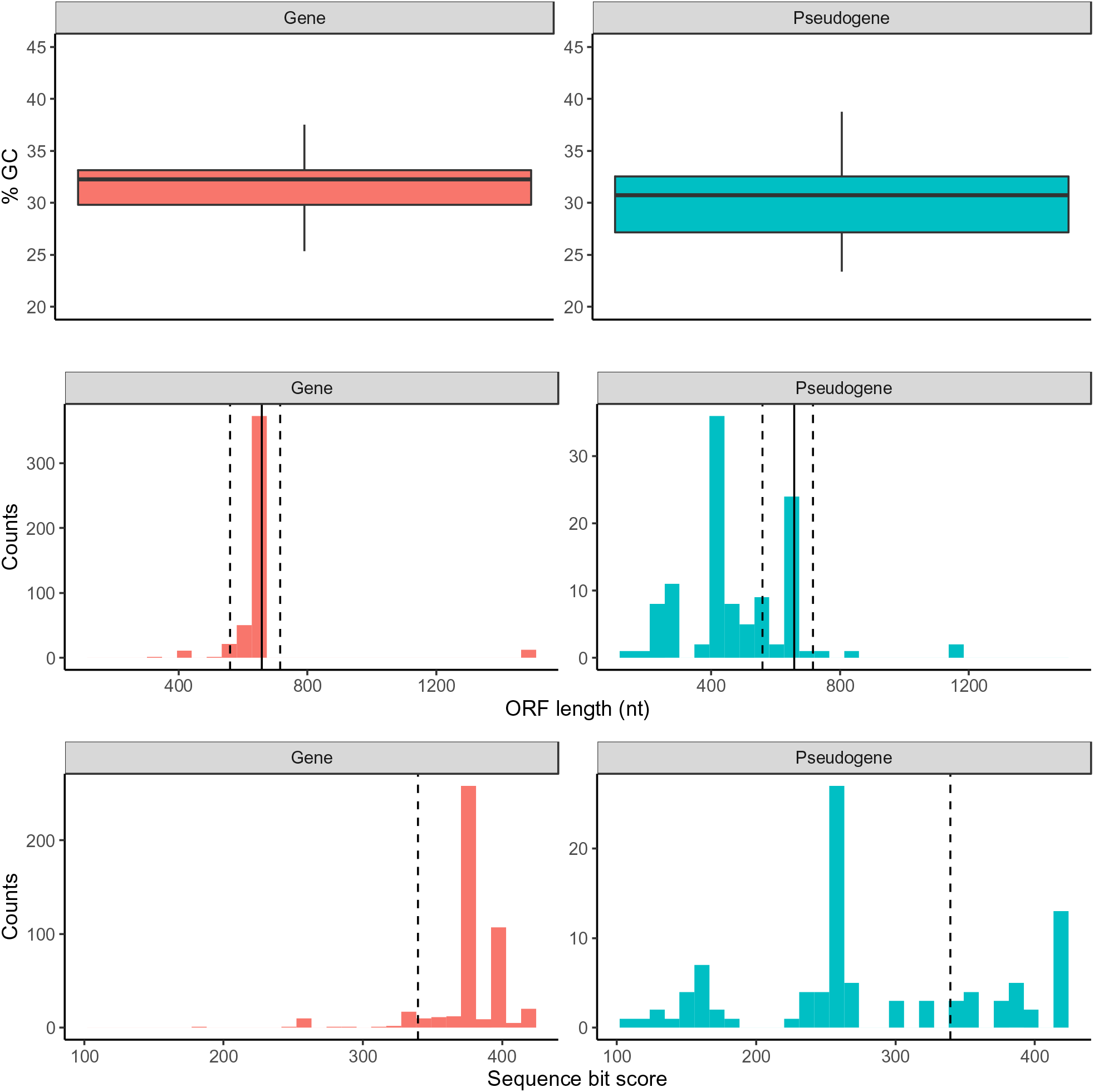
Arthropod COI pseudogenes tend to have lower GC content, shorter open reading frames, and smaller sequence bit scores. Based on the simulated DNA barcoding dataset described in Table 1. The top panel shows GC content (%) in gene and pseudogene sequences. The middle panel shows the sequence length distribution for the longest retained open reading frame. The solid vertical line indicates the length of a typical COI barcode at 658 bp. The two vertical dashed lines shows the boundaries for identifying ORFs with outlier lengths. The bottom panel shows the sequence bit score distribution after searching our sequences against a COI arthropod nucleotide profile hidden Markov model. The vertical dashed line shows the boundary for identifying small outlier scores.

Raw paired-end reads are merged using SEQPREP v1.3.2 [50]. This step looks for a minimum Phred quality score of 20 in the overlap region and requires a minimum 25 bp overlap. Primers are trimmed in two steps using CUTADAPT v2.6 requiring a Phred quality score of 20 at the ends to count matches/mismatches, no more than 3 Ns are allowed, and trimmed reads need to be at least 150 bp [51]. Sequence files are combined for a global analysis. Reads are dereplicated using VSEARCH v2.14.1 [52]. Denoised exact sequence variants (ESVs) are also generated using VSEARCH using the unoise3 algorithm [53]. This step clusters reads by 100% sequence identity, removes sequences with predicted errors, and globally rare sequences. Here we define rare sequences as clusters containing only one or two sequences. Putative chimeric sequences are removed using the uchime3_denovo algorithm in VSEARCH [54]. Denoised ORFs (ESVs) are taxonomically assigned using a naive Bayesian classifier trained with a COI reference set comprised of sequences mined from GenBank and the BOLD data releases [55, 56]. Rare sequences clusters are removed from each sample before printing the final file.

We used the pipeline with the two different pseudogene removal methods described in Part A. We then modified the pipeline to skip over several steps, one at a time, to see how this would affect the removal of apparent pseudogenes using the ORFfinder + profile HMM method: rare sequence removal, noise removal, chimeric sequence removal.

## Results

Our DNA barcode simulation that included 10 species with both gene and pseudogene sequences allowed us to compare differences in GC content, length, and dN/dS ratios. In Figure 2, we show that COI pseudogenes tend to have a slightly lower median GC content, shorter ORF lengths, and shorter full sequence bit score values in HMM profile analyses. Figure S1 shows how COI genes tend to accumulate substitutions in synonymous sites where a nucleotide changes does not result in the change of an amino acid; whereas COI pseudogenes tend to accumulate substitutions in non-synonymous sites where a nucleotide change results in the change of an amino acid. After correcting for pairwise comparisons that could yield unreliable dN/dS ratios, where the number of substitutions at synonymous sites is < 0.01 or > 2, we were only able to calculate dN/dS for COI gene sequences but not for pseudogene sequences. Due to the length variation in COI pseudogenes and their resulting ORFs it was difficult to obtain reliable codon alignments for dN/dS analysis. This method may be more suitable for detecting cryptic pseudogenes that have open reading frame lengths similar to functional COI ORFs. Top BLAST hit analysis shows that all pseudogenes had a top BLAST hit to another sequence from the expected species (92% - 100% identity). In some cases, the top BLAST match for a known pseudogene was to another COI sequence annotated as a nuclear copy of a mitochondrial gene. More often, the top match for a pseudogene was to a COI gene sequence. This indicates that in some cases, careful analysis of top BLAST hit output could help flag putative pseudogenes. Figures S2-S11 show COI phylograms for each species. In some cases, pseudogenes form their own clusters (ex. *Bemisia tabaci*, *Goneplax rhomboides*), often on long branches (ex. *Bemisia tabaci*, *Xylosandrus germanus*, *Triatoma dimidiate*, *Trialeurodes vaporariorum*, *Goneplax rhomboides*, *Ectatomma gibbum*), but occasionally pseudogenes are found in clades intermixed with regular genes and little sequence divergence to distinguish them (ex. *Melissotarsus insularis*, *Lepidocyrtus cyaneus*, *Halictus rubicundus*, *Cyphoderris monstrosa*).

Table 2 compares the sensitivity and specificity of two pseudogene removal methods on this dataset. Figure S12 shows how we calculated sensitivity and specificity for each pseudogene removal method. Sensitivity refers to the true positive rate, in this case the number of pseudogenes correctly filtered out of the dataset. Specificity refers to the true negative rate, in this case, the number of genes correctly retained. For our DNA barcoding simulated dataset including COI gene and pseudogene sequences from 10 species, sensitivity (73%) is slightly higher for the ORFfinder + HMM profile analysis pseudogene removal method and the specificity is the same for each pseudogene removal method (90%).

**Table 2.** Sensitivity and specificity for two pseudogene filtering methods. We include results from two approaches: Part A) We used a simulated DNA barcoding dataset with COI gene and pseudogene sequences from 10 species, Part B) we simulated pseudogenes from 100,000 BOLD COI sequences. To simulate pseudogenes, we either decreased the %GC content or introduced indels. Sensitivity refers to the true positive rate, our ability to correctly identify known or simulated pseudogenes. Specificity refers to the true negative rate, our ability to correctly identify real COI sequences (not pseudogenes).

| Experiment | Dataset | Type of mutations introduced | Sensitivity (%)<br>ORFfinder | ORFfinder + profile HMM analysis | Specificity (%)<br>ORFfinder | ORFfinder + profile HMM analysis |
| --- | --- | --- | --- | --- | --- | --- |
| Simulated DNA barcoding dataset. COI genes and pseudogenes from 10 species | Full length COI barcode and pseudogene sequences | N/A | 70 | 73 | 90 | 90 |
| Simulated metabarcode dataset | Full length COI barcode and simulated pseudogenes | GC content reduced | 31 | 27 | 99 | ~100 |
| Simulated metabarcode dataset | Full length COI barcode and simulated pseudogenes | Introduced indels | 88 | 94 | ~100 | ~100 |
| Simulated metabarcode dataset | Short COI barcode and simulated pseudogenes | GC content reduced | 17** - 50* | 6** - 15* | 99 | ~100 |
| Simulated metabarcode dataset | Short COI barcode and simulated pseudogenes | Introduced indels | 42** - 58* | 61** - 87* | 99 | 99* - ~100** |
| Simulated metabarcode dataset | Full length COI sequences and twice as many pseudogenes | GC content reduced | 17 | 0 | 99 | ~100 |
| Simulated metabarcode dataset | Full length COI sequences and twice as many pseudogenes | Introduced indels | 0 | 0 | ~100 | ~100 |
| Simulated metabarcode dataset | Full length COI sequences and half as many pseudogenes | GC content reduced | 39 | 36 | 95 | 96 |
| Simulated metabarcode dataset | Full length COI sequences and half as many pseudogenes | Introduced indels | 95 | 98 | 96 | 99 |

\* 5' fragment
\*\* 3' fragment

We used our observations from the simulated DNA barcode dataset with COI genes and pseudogenes from the same 10 species to guide the creation of a mock community comprised of 100,000 COI barcode sequences randomly sampled from BOLD where we could manipulate parameters in different ways. In our simulation study of full length COI sequences, we found that it was easier to filter out pseudogenes caused by increased indels (sensitivity 88-94%) rather than reduced GC content (sensitivity 27-31%) (Fig 3 and Table 2). As shown in Table 2, for full length COI barcode sequences, each pseudogene removal method performed with similar specificity (99-100%).

**Fig 3.**
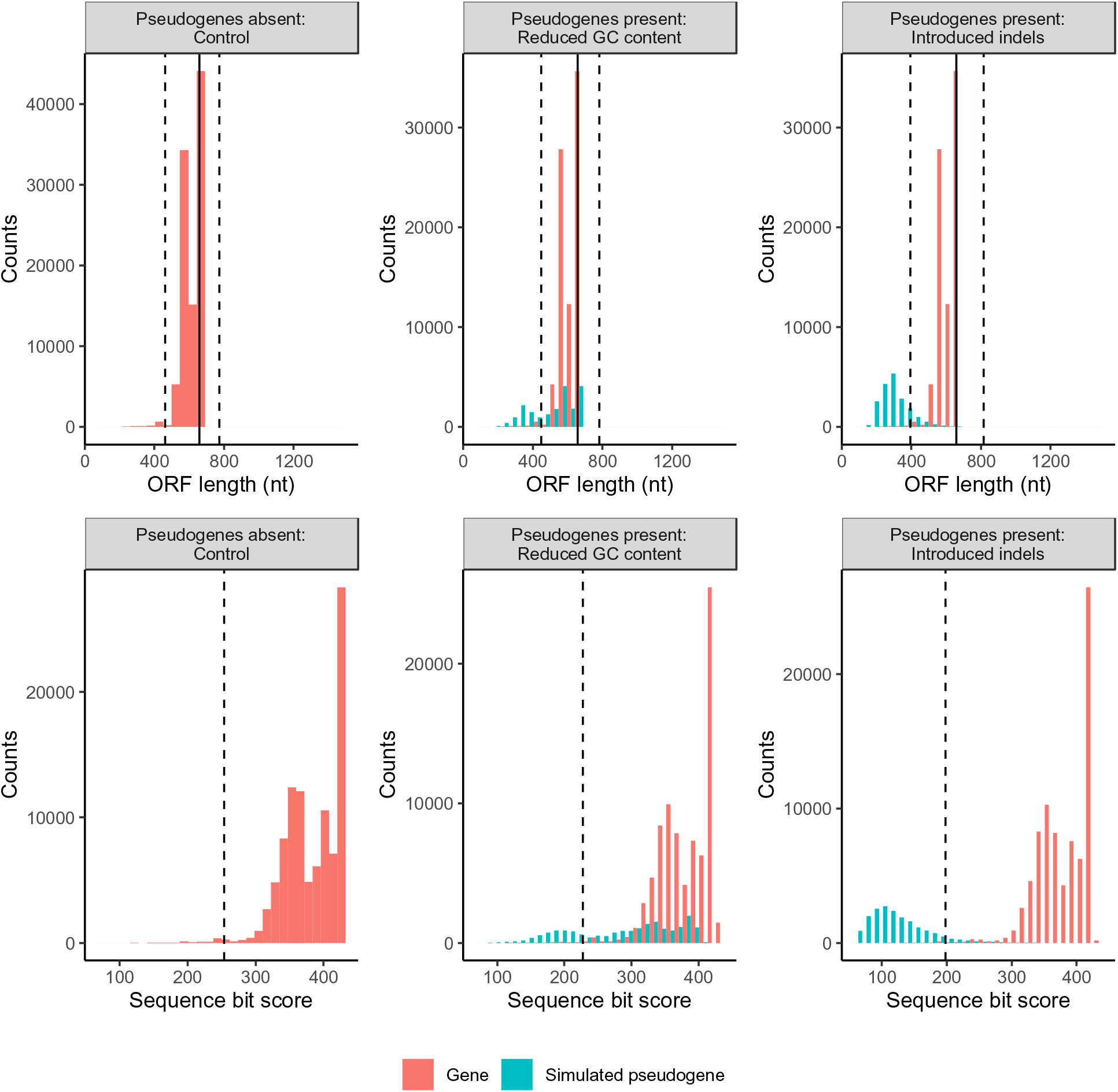
In a simulated mock arthropod community, reducing the GC content or introducing indels in COI sequences reduces ORF lengths and sequence bit scores. Each column shows the results from a particular simulation: a controlled community with pseudogenes absent, a community with pseudogenes that have a reduced GC content, and a community with pseudogenes where we have introduced indels. The top panel shows the length variation of sequences in the longest retained open reading frame. The solid vertical line indicates the length of a typical COI barcode at 658 bp. The two vertical dashed lines shows the boundaries for identifying ORFs with outlier lengths. The bottom panel shows the sequence bit score variation. The vertical dashed line shows the boundary for identifying sequences with low outlier scores.

We also performed additional simulations by adjusting the length of the COI barcodes from full length to half length (∼ 329 bp) as this is similar to the length of COI metabarcode sequences. As shown in Fig S13, it is more difficult to filter out short pseudogenes compared with full length COI barcodes. Table 2 shows that for half-length COI sequences, pseudogene removal sensitivity is better for pseudogenes generated by introducing indels (42-87%) rather than with pseudogenes where we reduced GC content (6-50%). Sensitivity is also generally higher when removing pseudogenes from the 5’ end of the COI barcode region (15-87%) compared with the 3’ end (6-61%). Pseudogene removal specificity is similar across pseudogene types and removal methods (99-100%).

Since we don’t really know how prevalent pseudogenes are in metabarcode datasets, we tested the effect of our pseudogene removal methods on a community where there are many pseudogenes (38% instead of 19% in previous analyses). Figure S14 shows that doubling the proportion of pseudogenes in the community greatly reduces the number of simulated pseudogenes removed with either method. As shown in Table 2, pseudogene removal sensitivity is poor (0-17%) but specificity is high using either removal method (99-100%). Next, we ran the opposite simulation where there are few pseudogenes in the community (9.5% instead of 19% in previous analyses). Figure S15 shows that reducing the number of pseudogenes in the community increases the number of simulated pseudogenes removed, especially when pseudogenes are caused by introducing indels. As Table 2 shows, the sensitivity of pseudogene removal is high when pseudogenes are created by introducing indels (95-98%), low when pseudogenes are created by reducing GC content (36-39%), and the specificity is high for any kind of simulated pseudogene or removal method (99-100%).

Because the ORFfinder + HMM profile analysis method for removing pseudogenes had the highest sensitivity for short COI metabarcodes when pseudogenes were simulated by introducing indels, we used this method to test our ability to remove pseudogenes with a real COI metabarcode dataset. Note that analyses were limited to only arthropod ESVs because most of the primer sets in the study were designed to specifically target this group in the original study (Table S2). As shown in Figure 4, the total number of arthropod ESVs was highest for the F230R amplicon (1,240) and least for the fwh1 amplicon (320). The greatest number of pseudogenes was detected and removed from the BR5 amplicon (19) and least for the ml-jg amplicon (1). Overall, the greatest percentage of pseudogenes out of all ESVs was detected from the BF2 amplicon (2.8%) and least for the ml-jg amplicon (0.1%). Because the F230R amplicon detected the greatest ESV richness, we used this amplicon to determine how existing bioinformatic processing steps affects pseudogene removal. Using the standard pipeline with ORFfinder + HMM profile analysis pseudogene removal, three F230R pseudogenes were removed from the dataset. Omitting the rare sequence removal step from the bioinformatic pipeline resulted in the largest number of pseudogenes detected, 34. Omitting the denoising step results in 1 pseudogene detected. Omitting the chimera removal step results in 16 pseudogenes removed. This suggests to us that at least some apparent pseudogenes are probably already being removed during regular bioinformatic processing, especially during the rare sequence removal step as we would expect from the literature [53, 54, 57–59].

**Fig 4.**
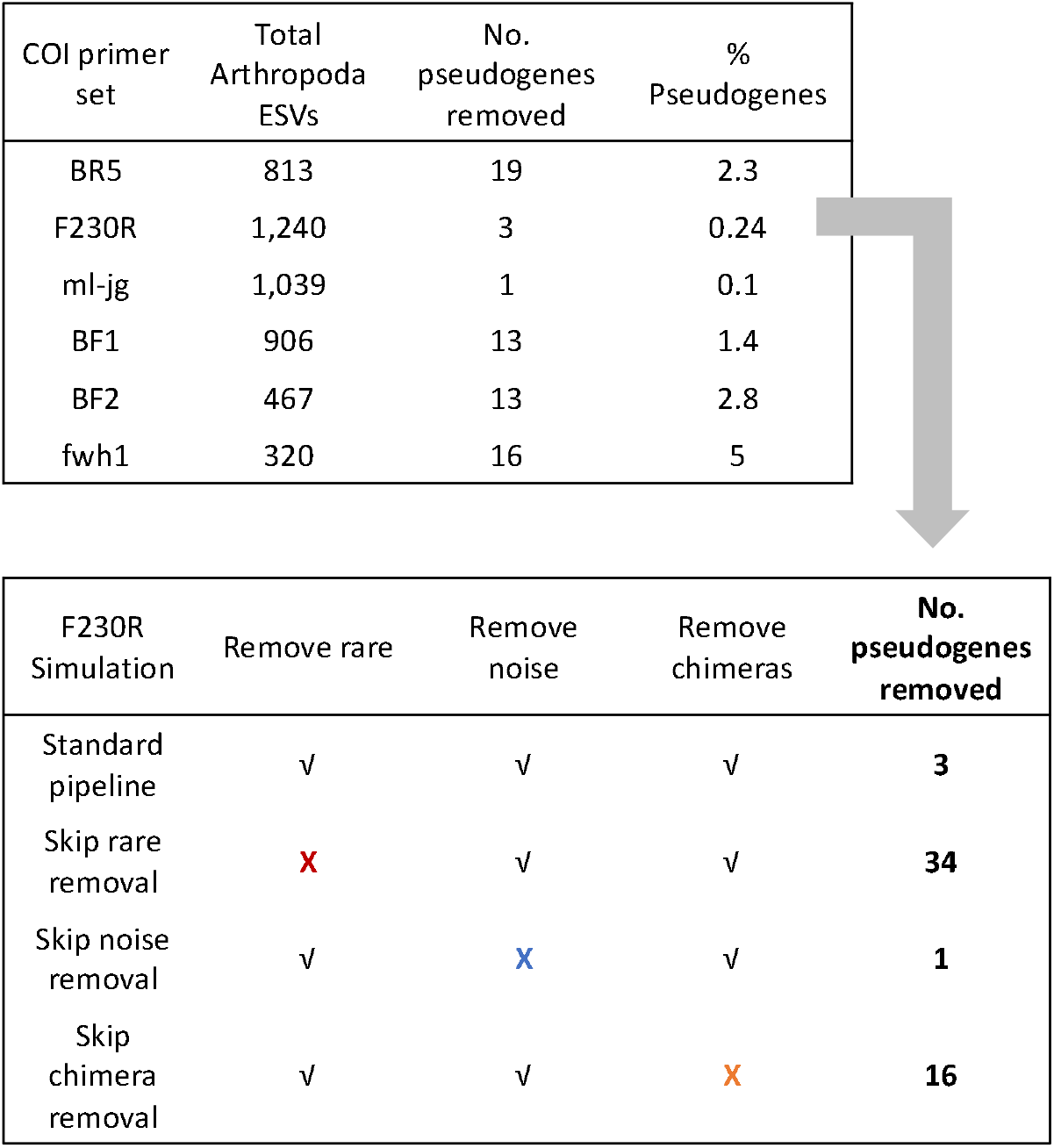
Removing rare sequences also removes apparent pseudogenes. The number of removed putative pseudogenes was calculated for each of the 5 amplicons from a real freshwater COI metabarcode dataset. Note, that we only compared results across Arthropoda ESVs. Using the standard bioinformatic pipline, the F230R amplicon recovered the greatest ESV richness (top box) so we used this as a test case for further simulations (bottom box). To determine whether current bioinformatic processing steps already help to remove apparent pseudogenes, we dropped one step at a time: removal of rare sequences, removal of noisy sequences, and removal of chimeric sequences.

## Discussion

Are all the COI sequences filtered out using ORFfinder + HMM profile analysis nuMTS? This method of sequence removal cannot distinguish between genuine pseudogenes and technical issues involving PCR or sequencing that causes indels, frameshifts, or the introduction of premature stop codons. It is possible that even after bioinformatic processing, artefactual sequences may be missed and subsequently be removed with these pseudogene removal methods. Although it is possible that genuine COI sequences could be removed using these methods, the specificity for pseudogenes is high (96-100%) and the number of COI gene sequences removed is very low in our simulated DNA barcode and metabarcode datasets.

There are also biological reasons why genuine mitochondrial sequences may be misclassified as pseudogenes. For example, in bivalves, male and female lineages of mitochondria may lead to fully functional gene copies with divergent sequences [15, 60, 61]. Though this type of sequence could complicate for COI barcoding or phylogenetic analysis, this would not be filtered out by our methods because as functional COI genes they are not expected to have frame shifts or shorter length that our method uses to flag potential pseudogenes. There are also cases in the literature where as a cell ages oxidative stress damages DNA that is then repaired by enzymes with reduced activity [15, 62]. Unrepaired mutations including deletions, duplications, and point mutations can accumulate in aging cells. Since truncated mtDNA can be replicated faster than full length mtDNA, it is possible for partially deleted mtDNA to accumulate [63]. Similarly, damaged DNA caused by poor preservation could cause COI sequences with frameshifts or premature stop codons to look like pseudogenes. It is quite likely that COI sequences with indels that lead to frameshifts and premature stop codons will be filtered out using the pseudogene removal methods we describe here whether the changes are technical or biological in nature.

How can pseudogenes be avoided? Indicators for the presence of pseudogenes include extra bands after PCR, sequence ambiguities when comparing both strands, frameshift mutations, premature stop codons, and unexpected phylogenetic position [18]. Strategies for avoiding pseudogenes in single specimens may include using muscle tissue for DNA extraction as it is naturally enriched with mtDNA, purifying mitochondria before DNA extraction, by amplifying long stretches of mtDNA with PCR, or targeting RNA using reverse transcription PCR [14, 18]. Even when working with environmental DNA samples, however, it can be possible to apply some of these techniques to avoid pseudogenes. For example, mitochondrial enrichment from homogenized tissues is possible and could be applied to freshwater benthic collections or insects collected from traps [64]. Additionally, long range PCR targeting mitochondrial DNA from water samples allowed for the construction of whole mitogenomes from fish [65]. Environmental RNA has also been used to detect microbes by targeting ribosomal RNA, this area has just begin to be explored using messenger RNA to target COI for metabarcoding [66–70]. For large scale studies, however, introducing additional steps such as mitochondrial purification or reverse transcription would be costly and time consuming.

Our results show that our ability to detect pseudogenes is hindered by short COI metabarcodes ∼ 300 bp in length or if the abundance of sequenced pseudogenes is very high. We show here that in a freshwater benthos COI metabarcode dataset, less than 3% of arthropod ESVs were removed as putative pseudogenes. It is quite possible that additional pseudogenes remain in the dataset, undetected by our pipeline. Our pseudogene removal methods cannot remove all pseudogenes, but remaining pseudogenes could still be useful for making higher level taxonomic assignments, though they may inflate richness at the species or haplotype level. Failure to remove low quality and artefactual sequences can result in inflated richness estimates in biodiversity studies, as has been shown for grashoppers and crayfish [14]. Pseudogenes are unlikely to affect community composition or beta diversity analyses if they are rare in the dataset as these analyses are less likely to be affected by the presence of rare sequences.

The use of phylogenetic based methods is common in COI barcoding studies, but the presence of pseudogenes could be a complication [14, 24, 26]. For example, a study of the great apes, showed that nuMTS are commonly sequenced in gorillas and complicate phylogenetic analyses [71]. It has also been suggested that pseudogenes are common in *Drosophila melanogaster* and in fish where they were once thought to be absent [72, 73]. The increasing use of COI metabarcodes for intraspecific analyses using ESVs could also be impacted by the presence of cryptic pseudogenes. The use of ORFfinder + HMM profile analysis, screening out hits with low outlier sequence bit scores, could be used as a first pass method for removing obvious pseudogenes. An automated method such as what we use in the SCVUC metabarcode pipelines in this study is more straight-forward to score compared with trying to identify pseudogenes from phylogenies by eye as branching patterns between genes and pseudogenes are not always clear cut. To detect cryptic pseudogenes careful analysis of species level sequence alignments should still be carried out to check for sequences with low GC content, high dN/dS ratios, indels, and codon usage bias.

Hidden Markov model profile analysis is not a commonly used method to process COI metabarcodes but it is used for many other applications. For example, the ITSx extractor is a program used to process fungal ITS metabarcodes by identifying and removing the conserved gene regions adjacent to the internal transcribed spacer regions (ITS1 and ITS2) [74]. HMMs are already used in the Pfam database of protein families [75]. HMM analysis is also used to place 16S rRNA gene sequences in a reference phylogeny in PICRUST2 [76]. The HMM profile analysis approach would be suitable for identifying gene sequences from protein coding markers such as rbcL and matK (plants), such that poor hits could be filtered out as putative pseudogenes. A multi-marker metabarcode pipeline that processes paired-end Illumina reads that provides a pseudogene filtering step for protein coding markers is the MetaWorks snakemake pipeline that can be found at https://github.com/terrimporter/MetaWorks. Furthermore, though our current work has focused on arthropod sequences, taxon-specific HMM profiles could be developed for additional macroinvertebrate groups of interest for biomonitoring such as tubellaria, gastropoda, bivalvia, polychaeta, oligochaeta, and hirudinea to permit more refined HMM-profile analyses [46]. It would also be useful to develop HMM profiles for other commonly used protein coding markers such as rbcL and matK to facilitate nuMT removal from large plant sequence datasets.

## Conclusions

We have shown that it is possible to screen out obvious pseudogenes using ORF length filtering alone or combined with HMM profile analysis for greater sensitivity when pseudogene sequences contain indels. Our pseudogenes removal approach was most effective on datasets of the full length COI barcode sequence region but is less effective for shorter sequences (∼ 300 bp). This is especially relevant now that newer sequencing technologies such as LoopSeq (compatible with Illumina sequencing platforms, but currently only available for RNA genes) or HiFi circular consensus sequencing (PacBio) could one day be used for COI metabarcoding targeting the full length of the barcoding region facilitating pseudogene detection [12, 77–79]. It would also be helpful if COI barcode studies reported and deposited full length verified pseudogenes into public databases when possible. Having key words such as ‘nuclear copy of mitochondrial gene’ or ‘pseudogene’ in the description would be essential to quickly flag hits to such sequences. As the analysis of metabarcode sequences from protein-coding genes shifts towards the use of exact sequence variants, it is more important than ever to reduce noise by removing pseudogenes when possible to avoid inflated richness estimates or misleading phylogenetic results. The incorporation of pseudogene filtering steps into widely used pipelines such is needed.

## List of abbreviations

BLAST: basic local alignment search tool
BOLD: Barcode of Life Data System
COI: cytochrome c oxidase subunit 1 gene
dN/dS: ratio of non-synonymous to synonymous substitions
ESV: exact sequence variant
GC content: guanine-cytosine content
HMM: Hidden Markov Model
ITS: internal transcribed spacer region in the ribosomal RNA operon
K2P: Kimura 2-parameter model of nucleotide substitution
matK: maturase K gene
mtDNA: mitochondrial DNA
nuMT: nuclear encoded mitochondrial sequence
NCBI: National Center for Biotechnology Information
ORF: open reading frame
OTU: operational taxonomic unit
rbcL: ribulose bisphosphate carboxylate large chain gene

## Declarations

### Ethics approval and consent to participate

Not applicable

### Consent for publication

Not applicable

### Availability of data and materials

All infiles and scripts used to parse data and generate figures are available from GitHub at xxx. The SCVUC COI metabarcode pipeline used in this study is also available on GitHub from https://github.com/Hajibabaei-Lab/SCVUC_COI_metabarcode_pipeline.

### Authors’ contributions

MH and TP conceived of the idea. TP conducted the analyses and wrote the manuscript. MH provided critical input into analysis methods and the manuscript. MH provided funding and computational resources. Both authors edited, read, and approved the final manuscript.

## Acknowledgements

We would like to thank the Hajibabaei group for their support and helpful discussions.

## Supplementary Material

**Table S1.** Description of the datasets analyzed in Part A and Part B.

| Experiment | Dataset | Proportion of dataset comprised of pseudogenes (%) | Average gene length (bp) | Average pseudogene length (bp) | Gene GC content (%) | Pseudogene GC content (%) |
| --- | --- | --- | --- | --- | --- | --- |
| Part A | Simulated DNA barcode dataset | 19 | 659.6 | 508.1 | 32.0 | 30.8 |
| Part B | Control mock community with 100,000 randomly sampled sequences from BOLD | 0 | 615 | NA | 31 | NA |
| Part B | Mock community with decreased GC content | 19 | 615 | 615 | 31 | 29 |
| Part B | Mock community with increased indels | 19 | 615 | 607 | 31 | 31 |
| Part B | Control mock community with half-length sequences | 0 | 307** - 308* | NA | 30*-32** | NA |
| Part B | Mock community with half-length sequences and decreased GC content | 19 | 307** - 308* | 308 | 30*-32** | 28-29 |
| Part B | Mock community with half-length sequences and increased indels | 19 | 307** - 308* | 304 | 30*-32** | 31-32 |
| Part B | Control mock community with twice 100,000 randomly sampled BOLD sequences | 0 | 622 | NA | 31 | NA |
| Part B | Mock community with twice the number of<br>pseudogenes<br>with decreased<br>GC content | 38 | 622 | 622 | 31 | 28 |
| Part B | Mock<br>community<br>with twice the<br>number of<br>pseudogenes<br>with increased<br>indels | 38 | 622 | 614 | 31 | 32 |
| Part B | Control mock<br>community of<br>100,000<br>randomly<br>sampled<br>sequences<br>from BOLD | 0 | 622 | NA | 31 | NA |
| Part B | Mock<br>community<br>with halved<br>number of<br>pseudogenes<br>with decreased<br>GC content | 9.5 | 622 | 623 | 31 | 28 |
| Part B | Mock<br>community<br>with halved<br>number of<br>pseudogenes<br>with increased<br>indels | 9.5 | 622 | 615 | 31 | 32 |

\* 5' fragment
\*\* 3' fragment

**Table S2.**
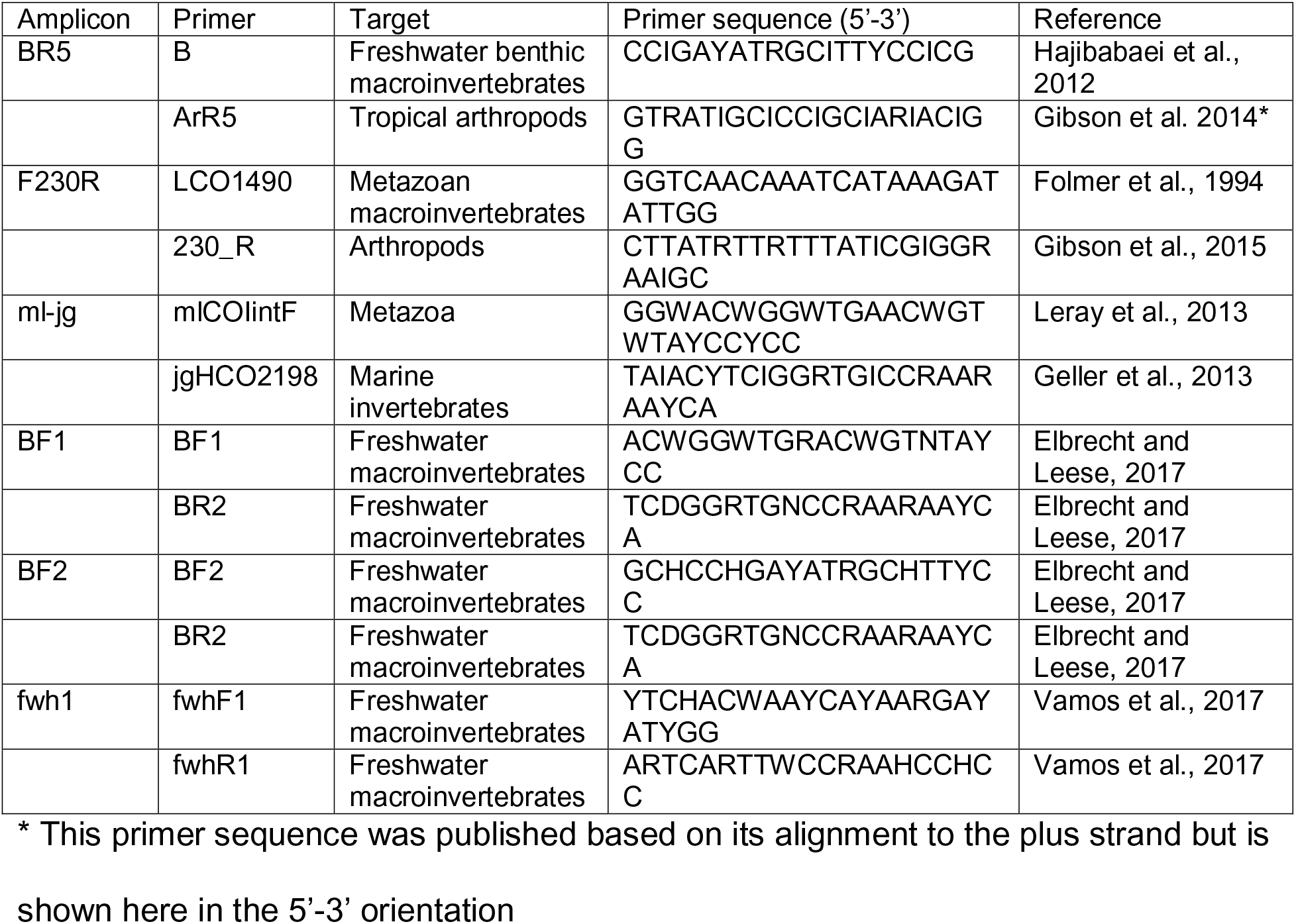
Primers used in the freshwater benthos COI metabarcode dataset used in Part C (Hajibabaei et al., 2019 PLoS ONE).

**Fig S1.**
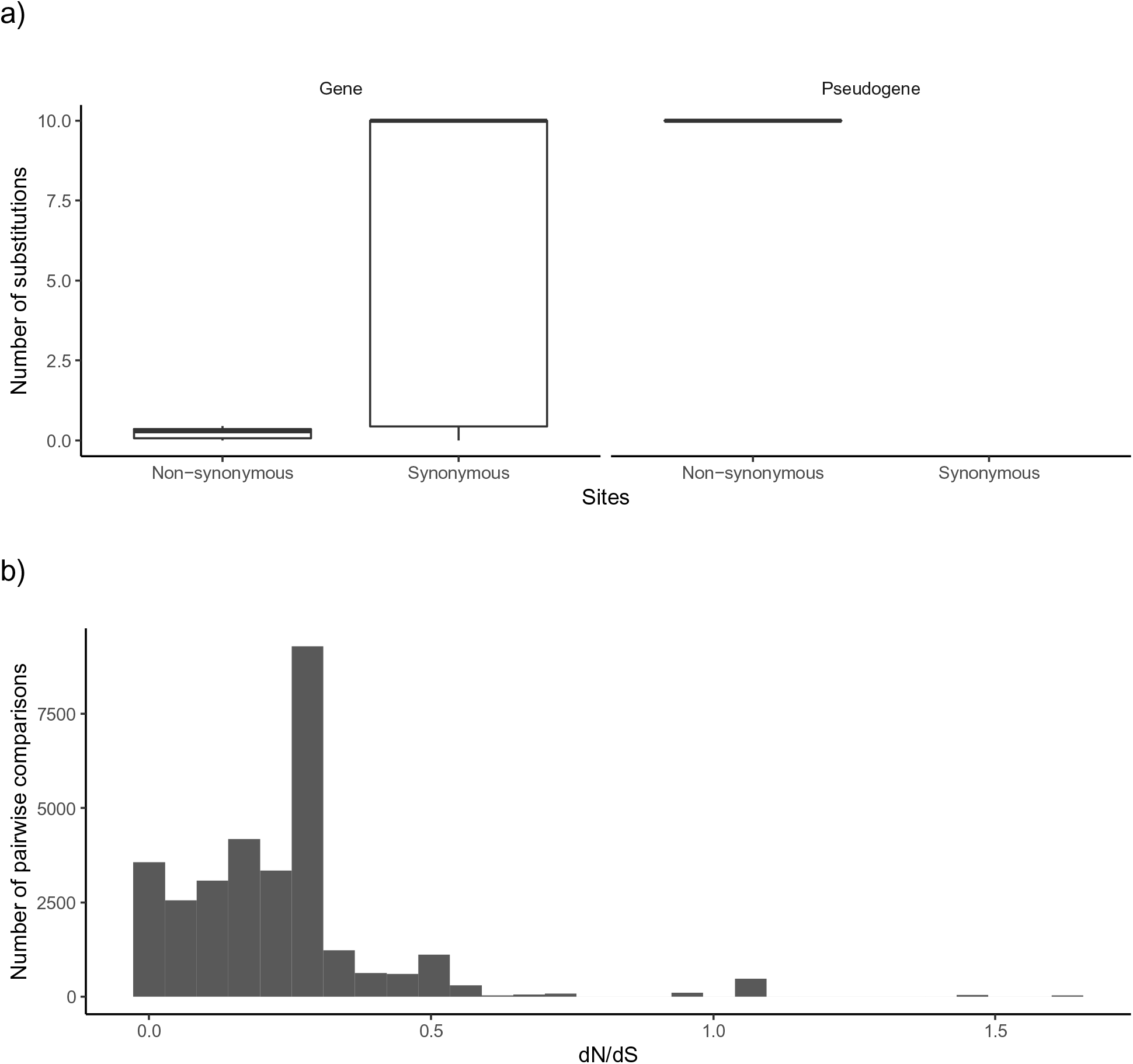
COI gene sequences accumulate substitutions in synonymous sites. For 10 species with annotated COI genes and pseudogenes, we did a pairwise comparison of nucleotide substitutions in non-synonymous and synonymous sites: a) COI barcode sequences tend to accumulate substitutions in synonymous sites. In contrast, COI pseudogenes tend do accumulate substitutions in non-synonymous sites. After filtering out pairwise comparisons between species with < 0.01 substitutions in synonymous sites (sequences too similar to yield a reliable dN/dS estimate) or > 2 substitutions in synonymous sites (sequences that have accumulated too many substitutions to yield a reliable dN/dS estimate), it was only possible to analyze dN/dS ratios for COI barcode sequences. b) Most pairwise comparisons of COI gene sequences resulted in dN/dS ratios < 1 consistent with purifying selection pressure and the conservation of a protein sequence.

**Fig S2.**
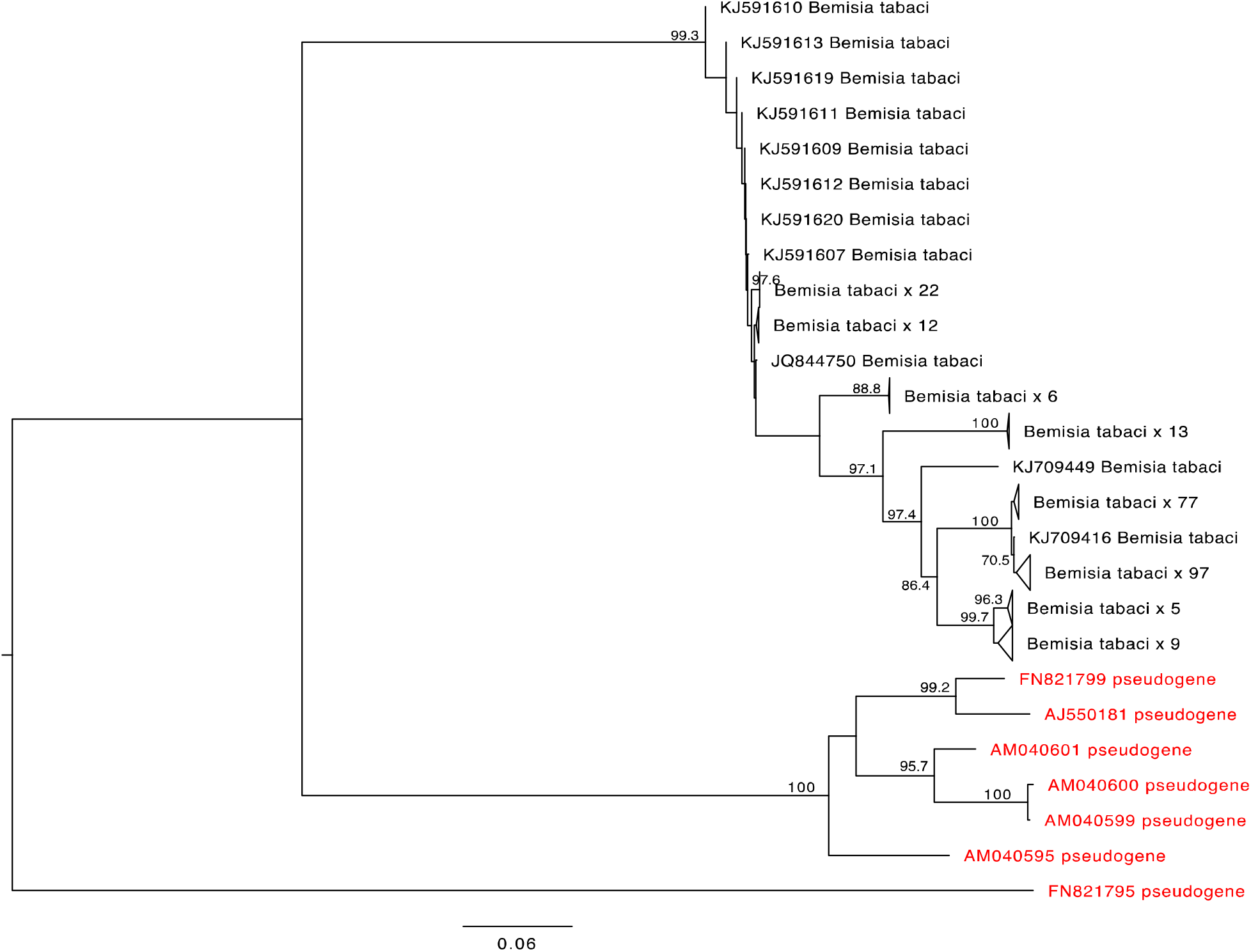
*Bemisia tabaci* COI pseudogenes cluster together on long branches. A mid-point rooted neighbor joining phylogram using the Kimura 2-parameter model of nucleotide substitution included gene and known pseudogene sequences. Sequences annotated in GenBank as a nuclear copy of a mitochondrial gene are shown in red. Nodes with greater than 70% bootstrap support are labelled.

**Fig S3.**
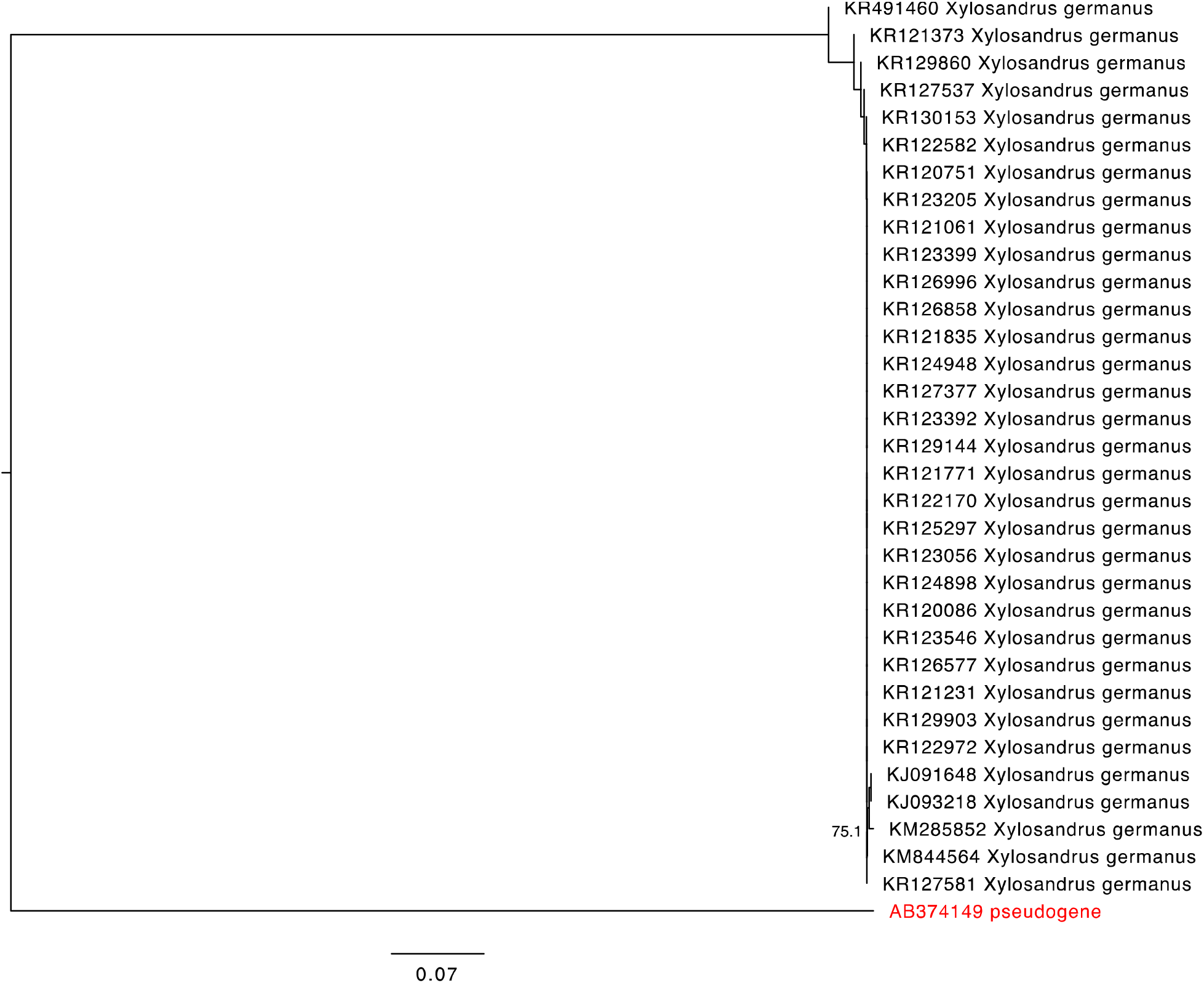
A single *Xylosandrus germanus* COI pseudogene sequence is found on a long branch. A mid-point rooted neighbor joining phylogram using the Kimura 2-parameter model of nucleotide substitution included COI gene sequences as well as a sequence annotated in GenBank as a nuclear copy of a mitochondrial gene (red). Nodes with greater than 70% bootstrap support are labelled.

**Fig S4.**
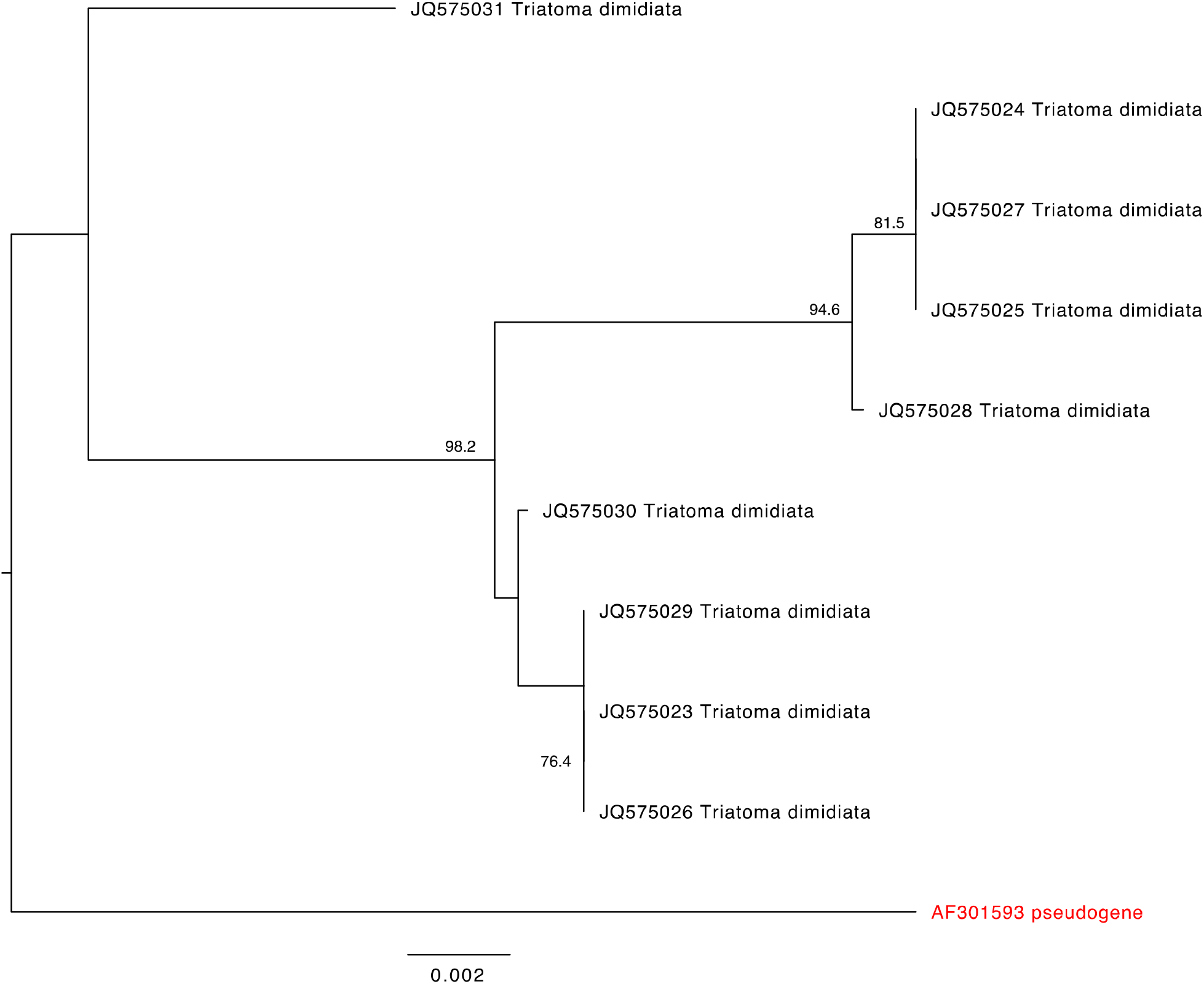
A single *Triatoma dimidiata* COI pseudogene sequence is found on a long branch. A mid-point rooted neighbor joining phylogram using the Kimura 2-parameter model of nucleotide substitution included COI gene sequences as well as a sequence annotated in GenBank as a nuclear copy of a mitochondrial gene (red). Nodes with greater than 70% bootstrap support are labelled.

**Fig S5.**
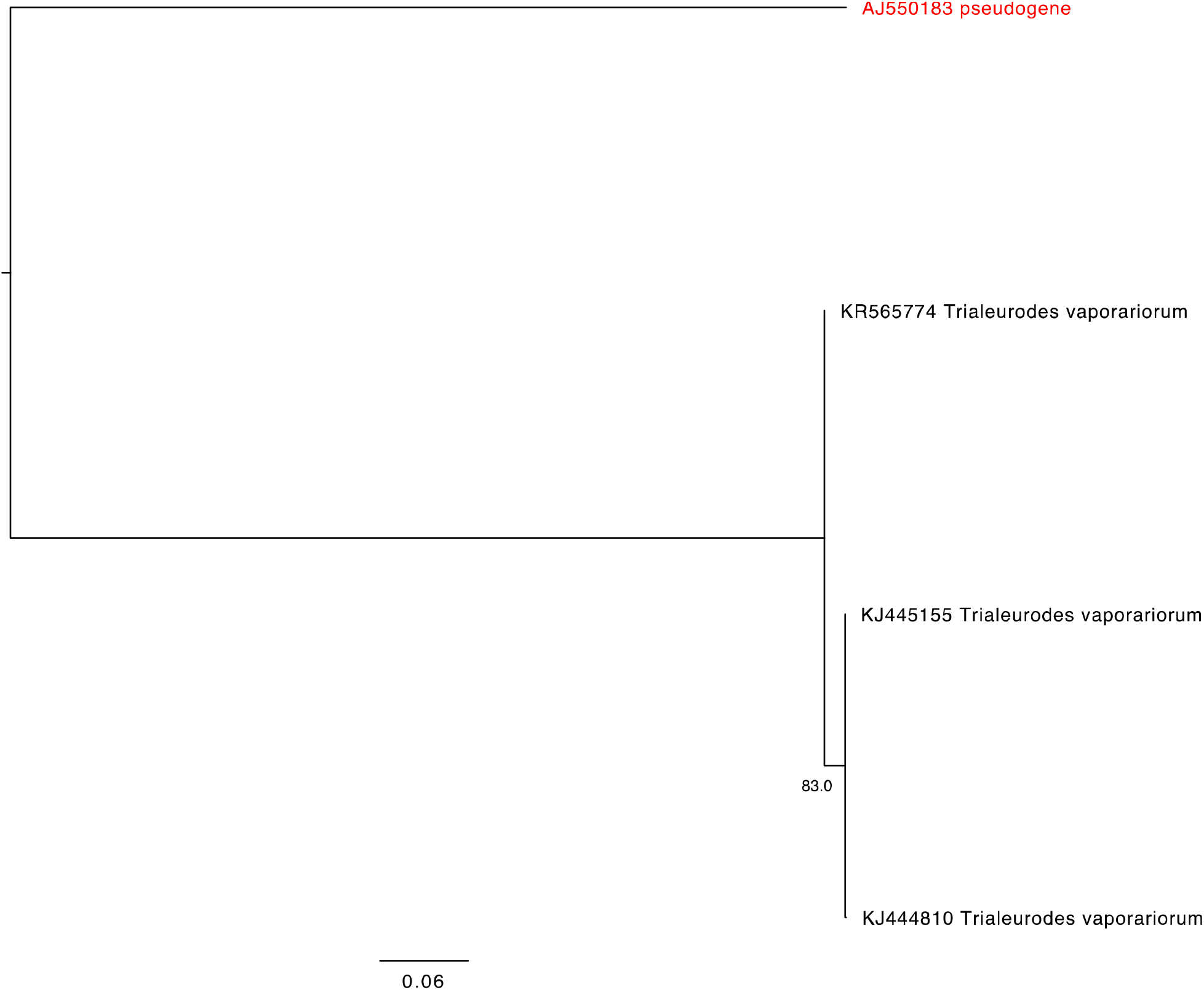
A single *Trialeurodes vaporariorum* COI pseudogene sequence is found on a long branch. A mid-point rooted neighbor joining phylogram using the Kimura 2-parameter model of nucleotide substitution included COI gene sequences as well as a sequence annotated in GenBank as a nuclear copy of a mitochondrial gene (red). Nodes with greater than 70% bootstrap support are labelled.

**Fig S6.**
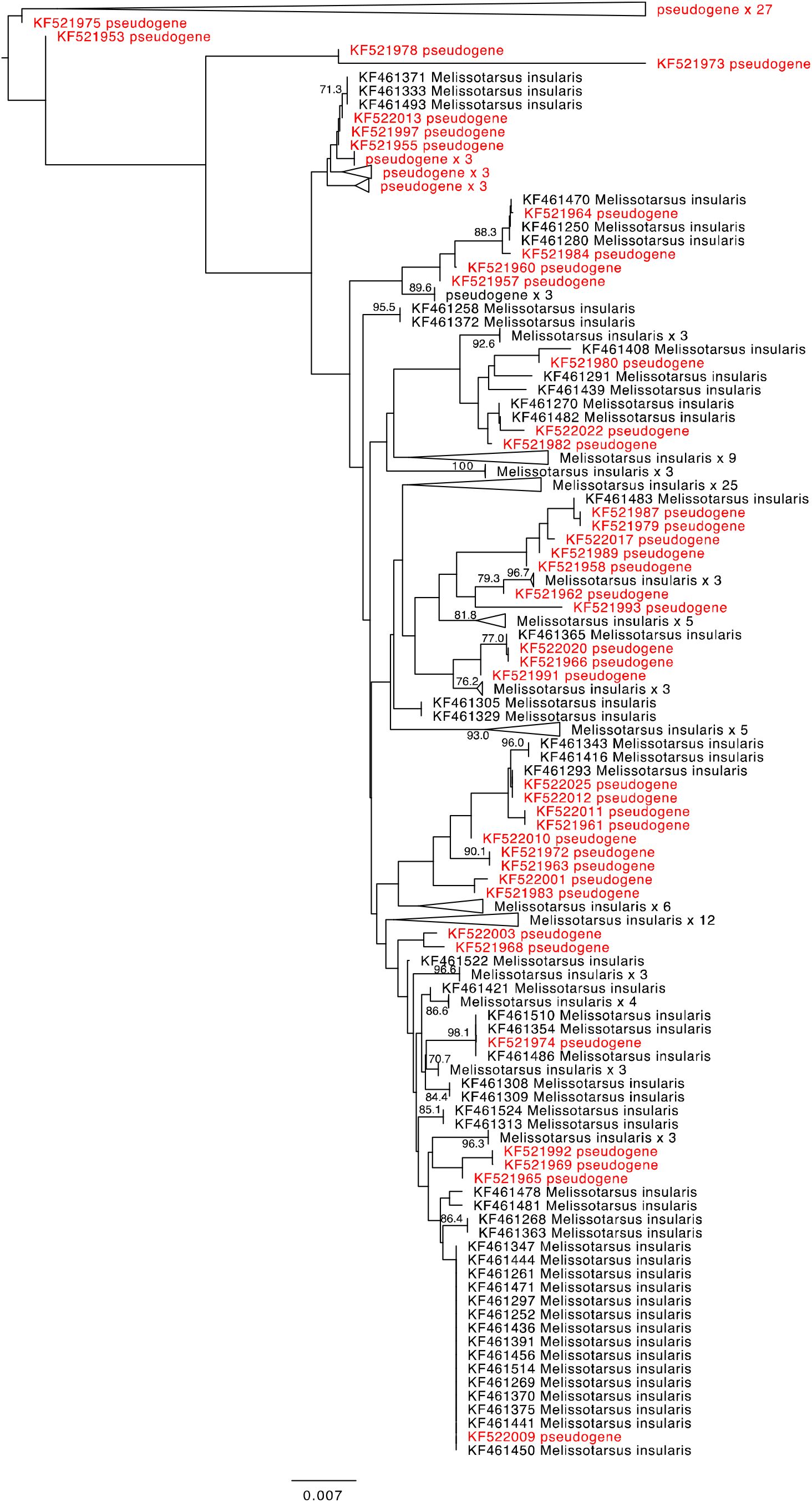
*Melissotarsus insularis* COI gene and annotated pseudogene sequences are often found in intermixed clusters. A mid-point rooted neighbor joining phylogram using the Kimura 2-parameter model of nucleotide substitution included COI gene sequences as well as sequences annotated in GenBank as a nuclear copy of a mitochondrial gene (red). Nodes with greater than 70% bootstrap support are labelled. Clusters of nearly identical sequences were collapsed.

**Fig S7.**
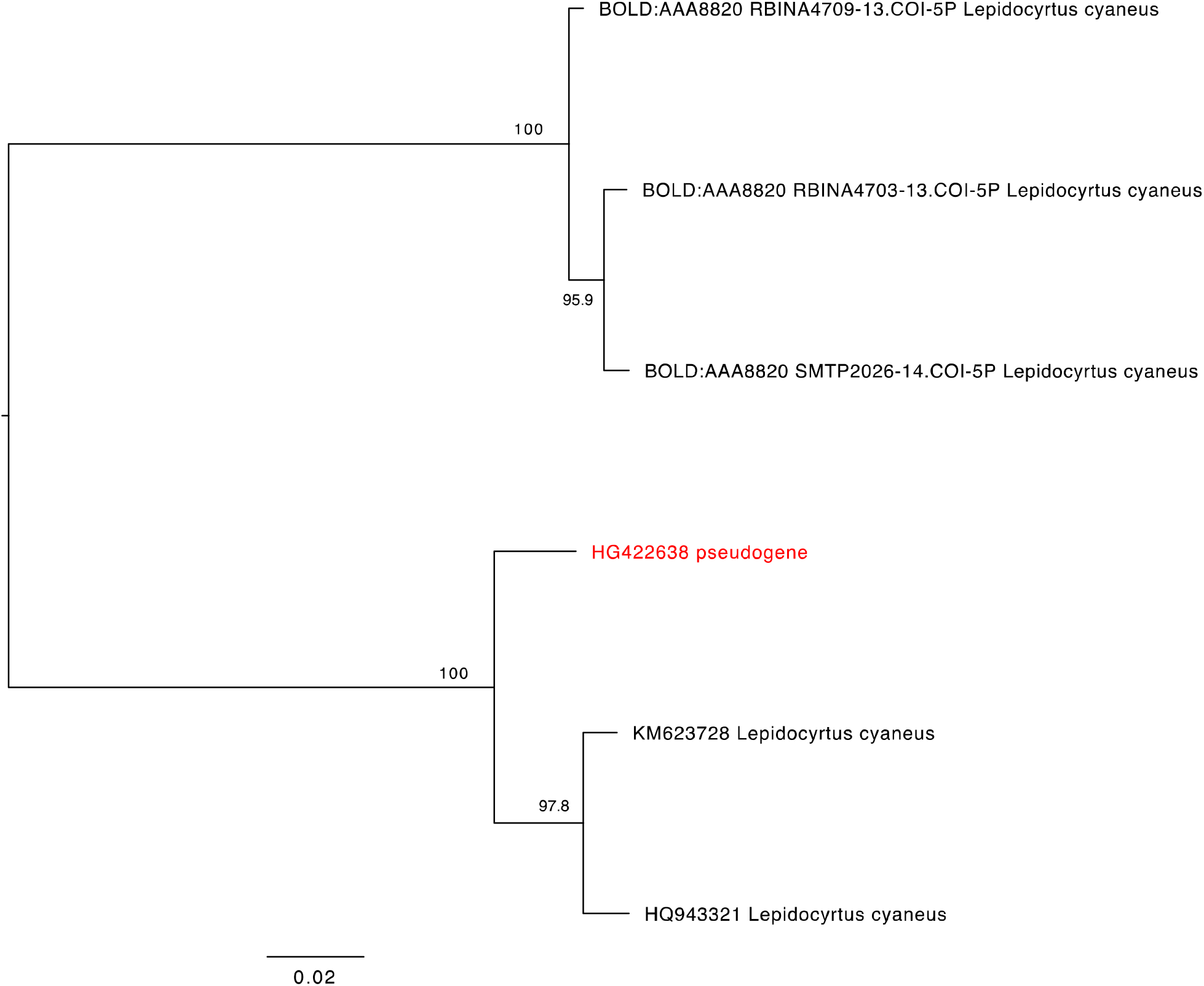
A single *Lepidocyrtus cyaneus* COI pseudogene sequence clusters with other gene sequences. A mid-point rooted neighbor joining phylogram using the Kimura 2-parameter model of nucleotide substitution included COI gene sequences as well as a sequence annotated in GenBank as a nuclear copy of a mitochondrial gene (red). Nodes with greater than 70% bootstrap support are labelled.

**Fig S8.**
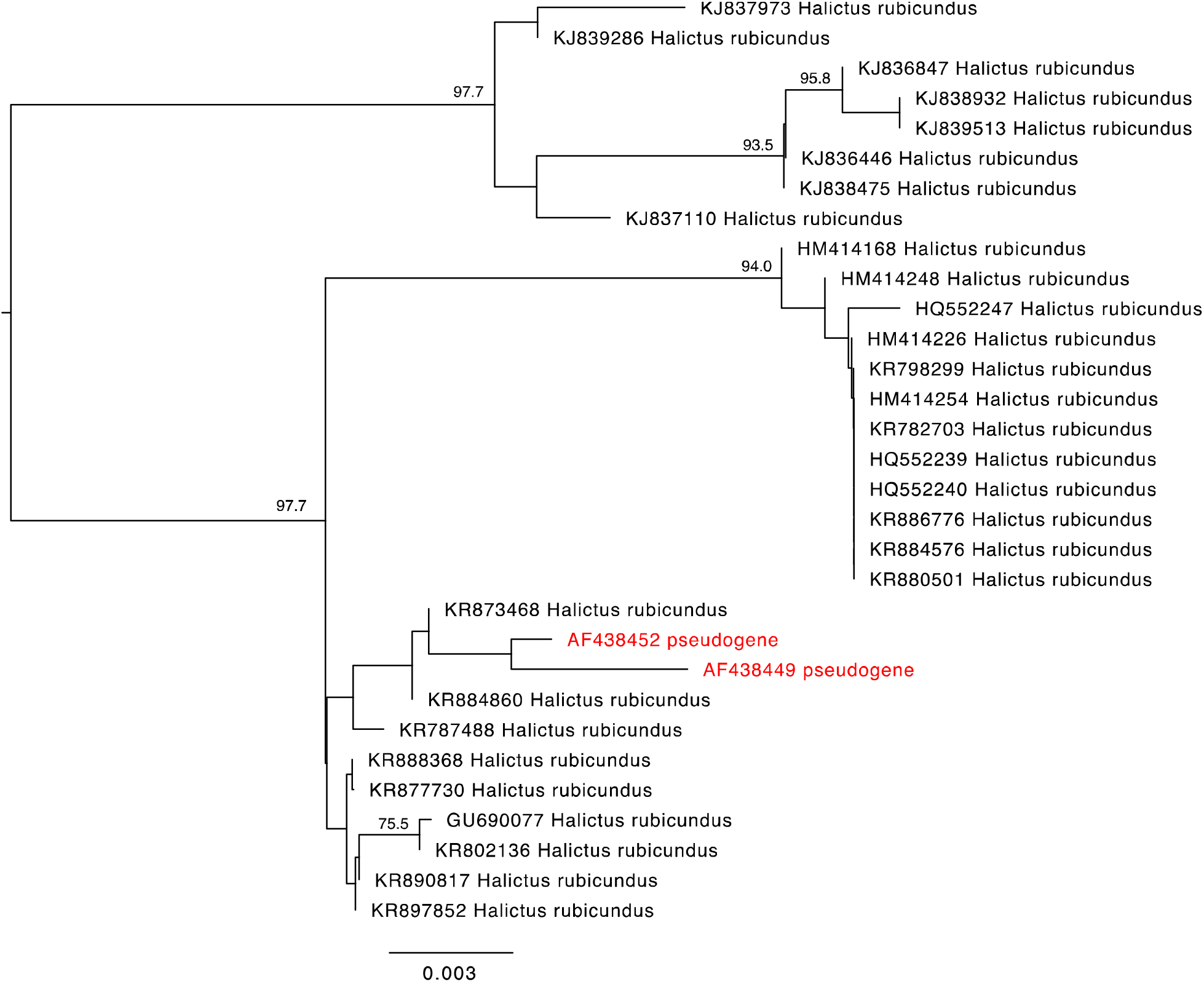
Two *Halictus rubicundus* COI pseudogene sequences cluster together near other gene sequences. A mid-point rooted neighbor joining phylogram using the Kimura 2-parameter model of nucleotide substitution included COI gene sequences as well as two sequences annotated in GenBank as a nuclear copy of a mitochondrial gene (red). Nodes with greater than 70% bootstrap support are labelled.

**Fig S9.**
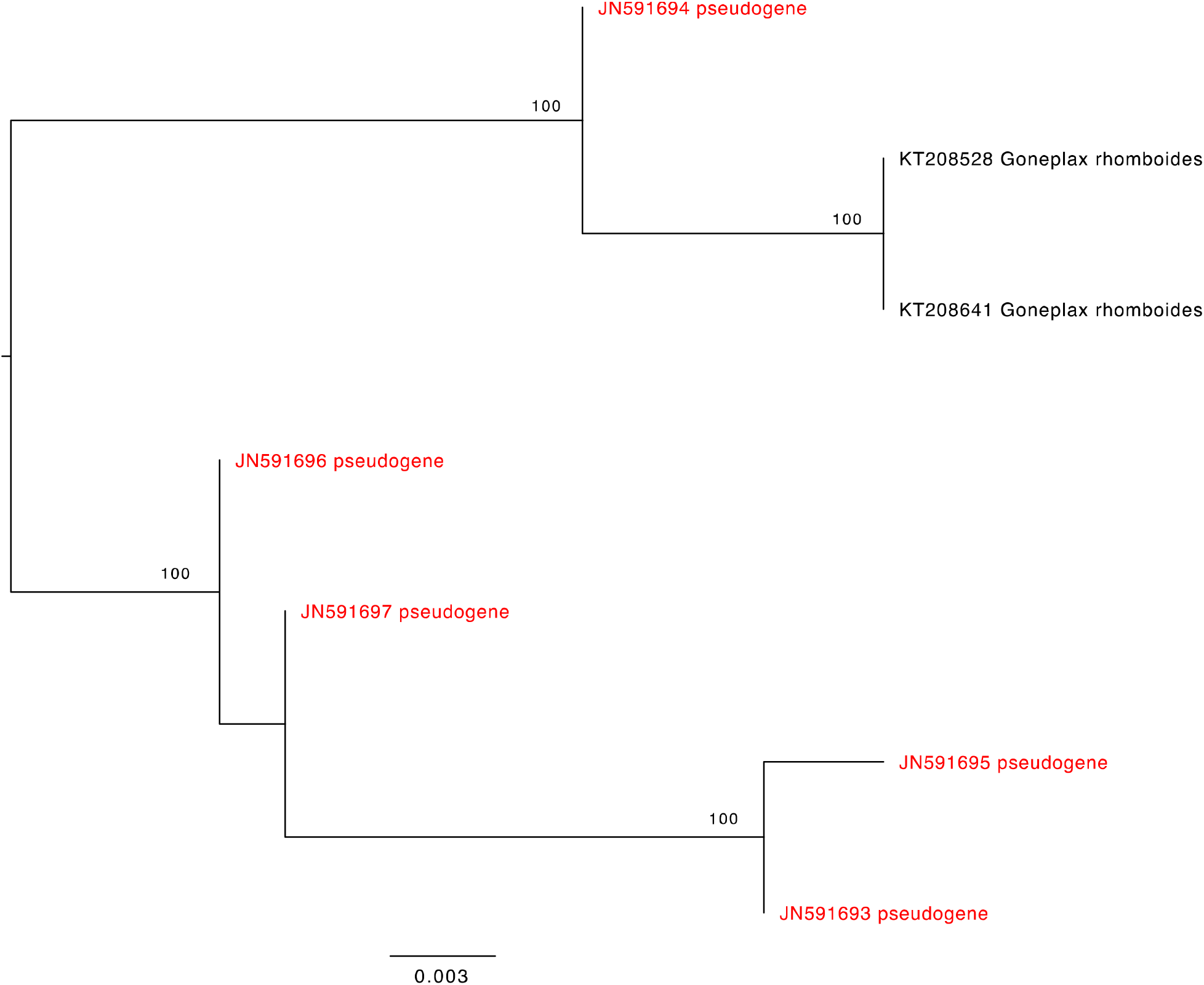
Several *Goneplax rhomboides* COI pseudogene sequences cluster together. A mid-point rooted neighbor joining phylogram using the Kimura 2-parameter model of nucleotide substitution included COI gene sequences as well as sequences annotated in GenBank as a nuclear copy of a mitochondrial gene (red). Nodes with greater than 70% bootstrap support are labelled.

**Fig S10.**
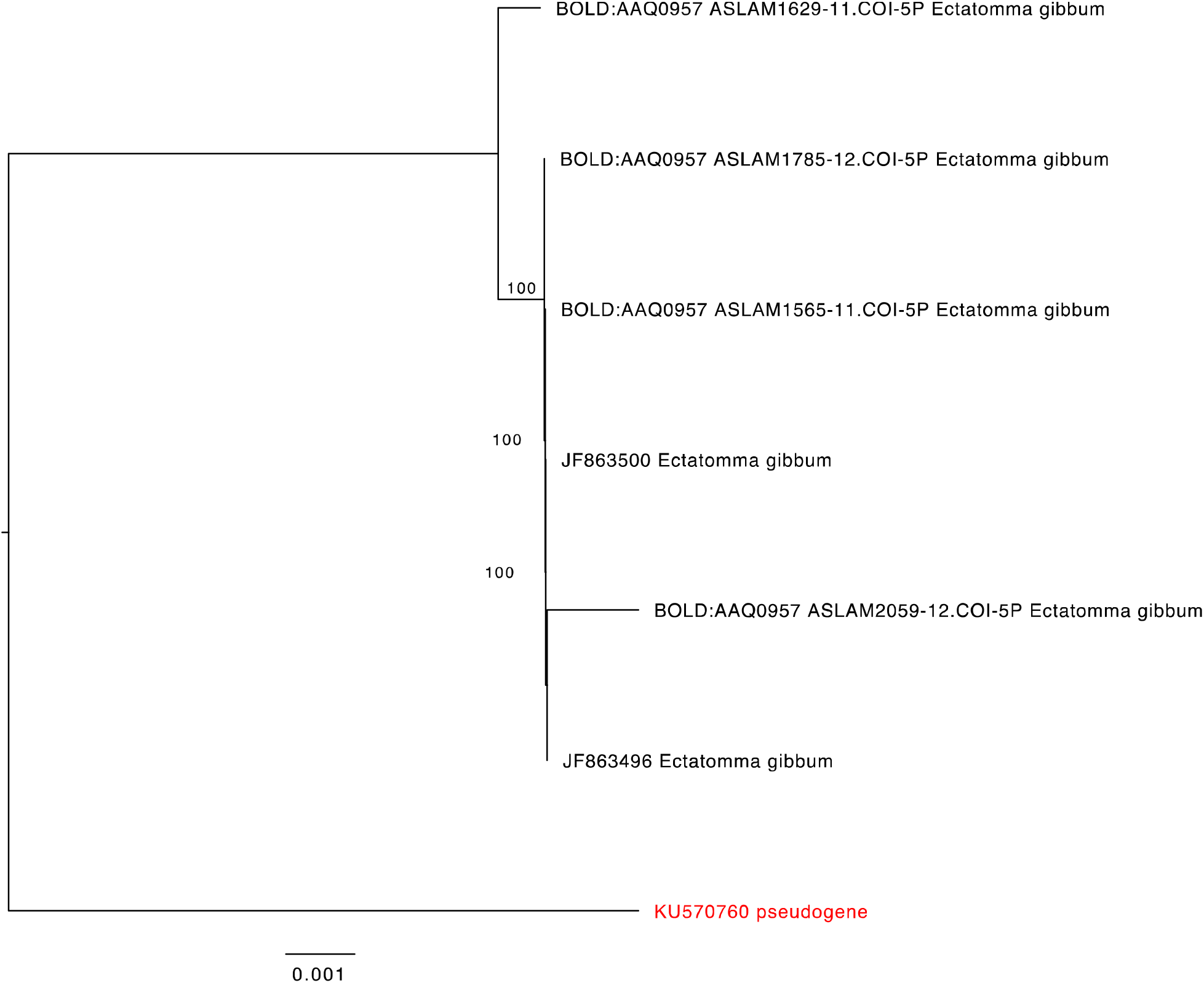
A single *Ectatomma gibbum* COI pseudogene sequence is found on its own branch. A mid-point rooted neighbor joining phylogram using the Kimura 2-parameter model of nucleotide substitution included COI gene sequences as well as a sequence annotated in GenBank as a nuclear copy of a mitochondrial gene (red). Nodes with greater than 70% bootstrap support are labelled.

**Fig S11.**
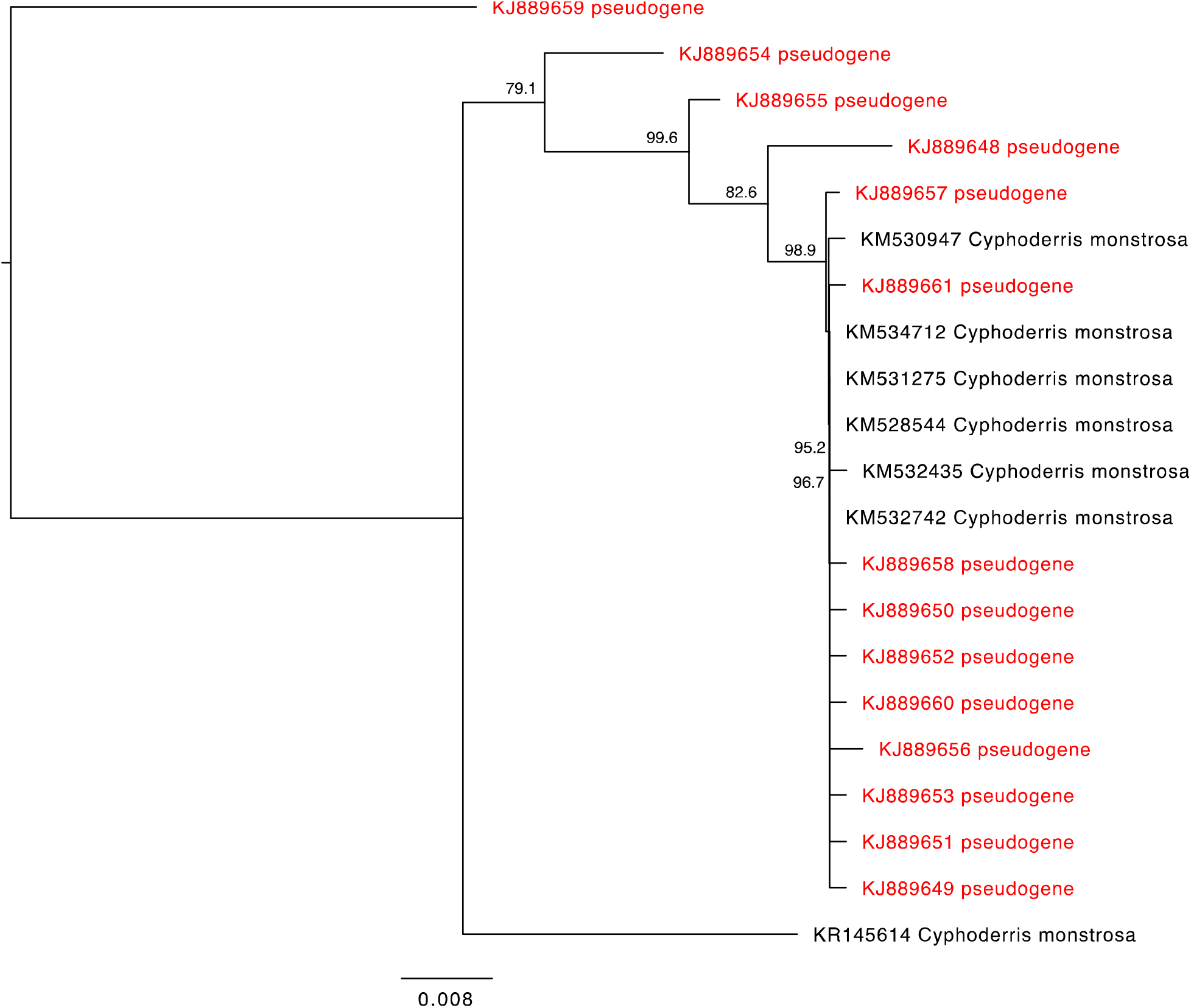
*Cyphoderris monstrosa* COI gene and annotated pseudogene sequences sometimes cluster with regular gene sequences. A mid-point rooted neighbor joining phylogram using the Kimura 2-parameter model of nucleotide substitution included COI gene sequences as well sequences annotated in GenBank as a nuclear copy of a mitochondrial gene (red). Nodes with greater than 70% bootstrap support are labelled.

**Fig S12.**
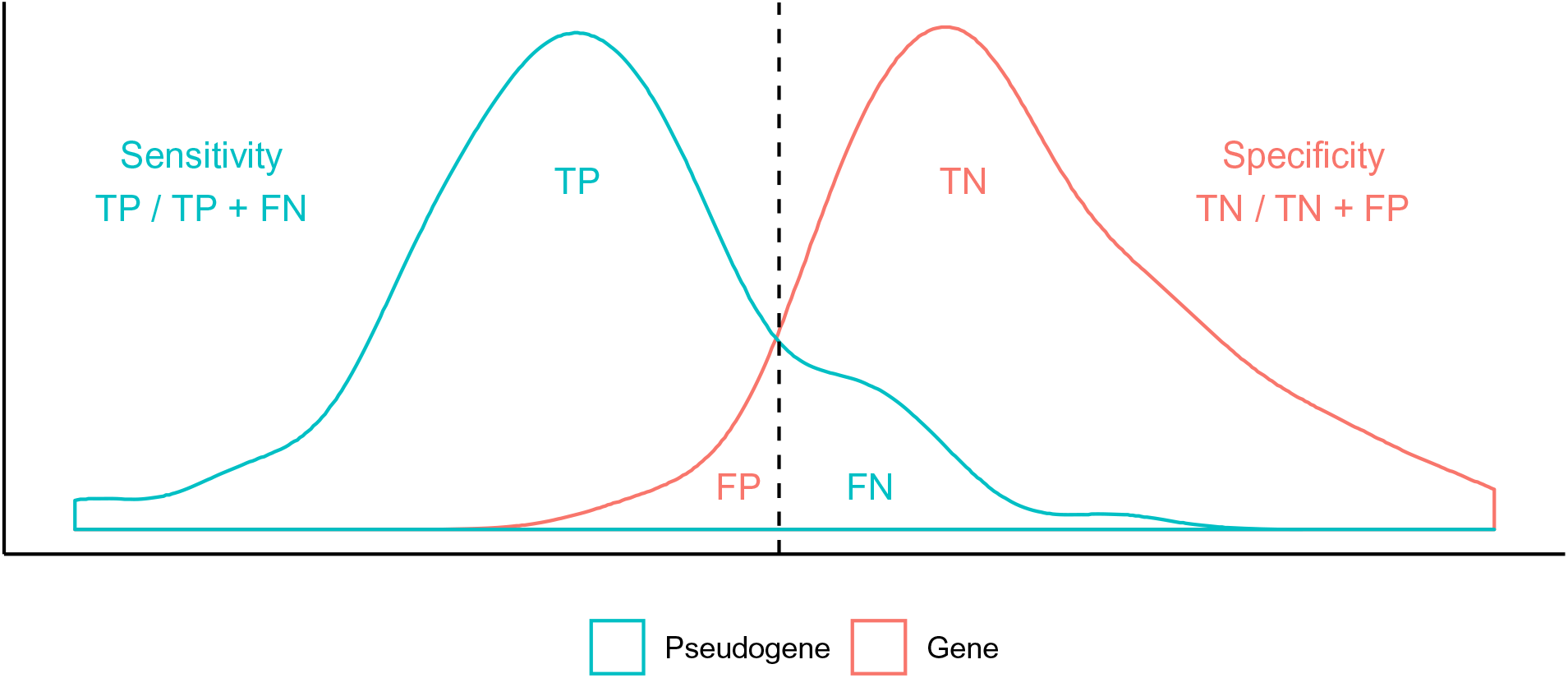
Sensitivity and specificity were used to assess the effectiveness of our two pseudogene filtering approaches. The vertical dashed line represents a threshold used to delimit COI pseudogene sequences. The ability to detect pseudogenes represents the positive condition. Correctly removed pseudogenes are true positives (TP). Incorrectly filtered COI gene sequences (genes) represents false positives (FP). Correctly retained genes represents true negatives (TN). Incorrectly retained pseudogenes represents false negatives (FN).

**Fig S13.**
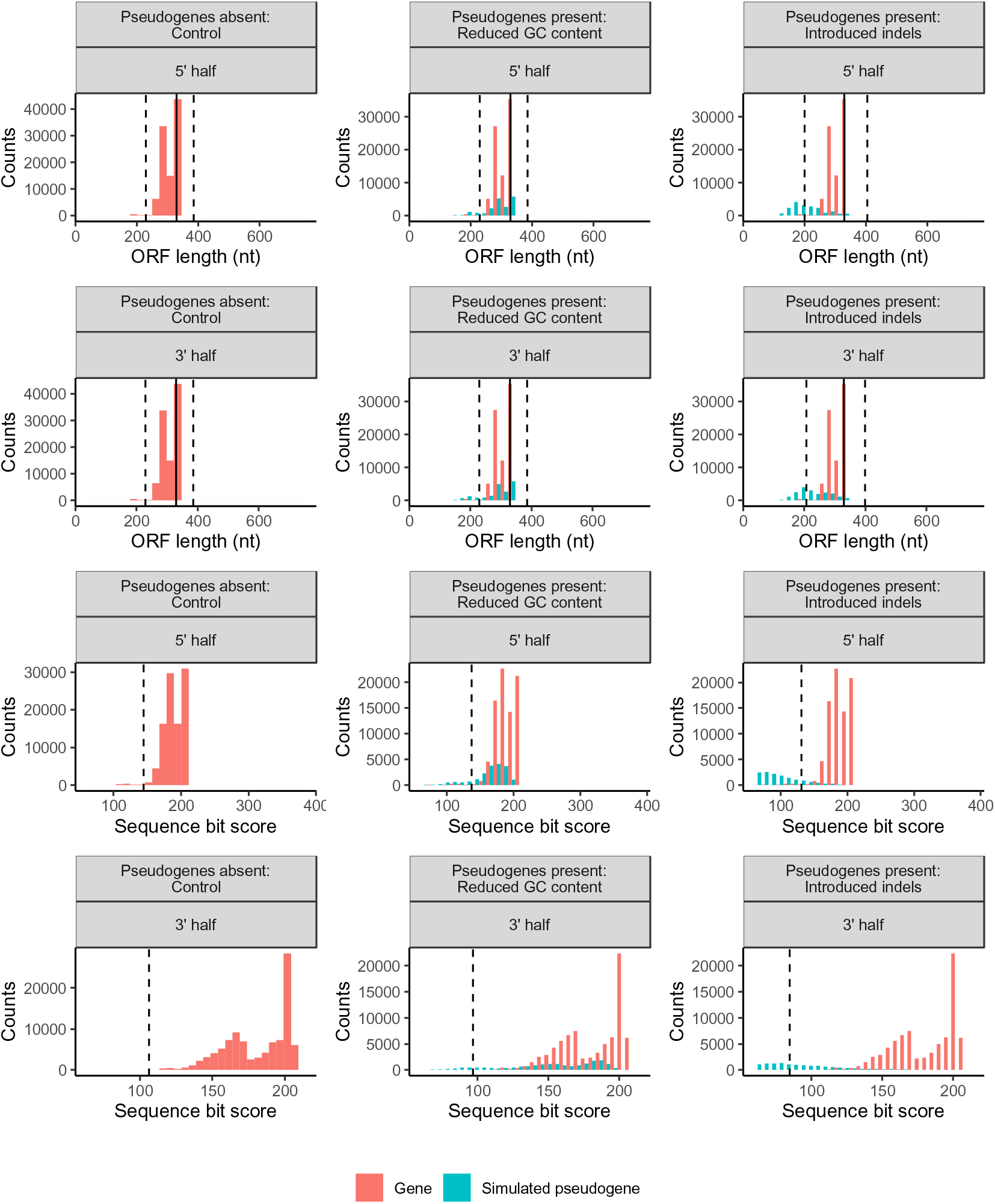
Halving COI sequence lengths results in fewer simulated pseudogenes removed compared with full length COI barcode sequences. Each column shows the results from a particular simulation: a controlled community with pseudogenes absent, a community with simulated pseudogenes with a reduced GC content, and a community with simulated pseudogenes with introduced indels. The top two panels show the length variation of sequences in the longest retained open reading frame for short sequences sampled from the 5’ and 3’ end of COI barcode sequences. The solid vertical line indicates half the length of a typical COI barcode at 329 bp. The two vertical dashed lines shows the boundaries for identifying ORFs with outlier lengths. The bottom two panels show the nucleotide bit score for short sequences sampled from the 5’ and 3’ ends of COI barcode sequences. The dashed vertical line shows the boundary for identifying sequences with unusually short scores.

**Fig S14.**
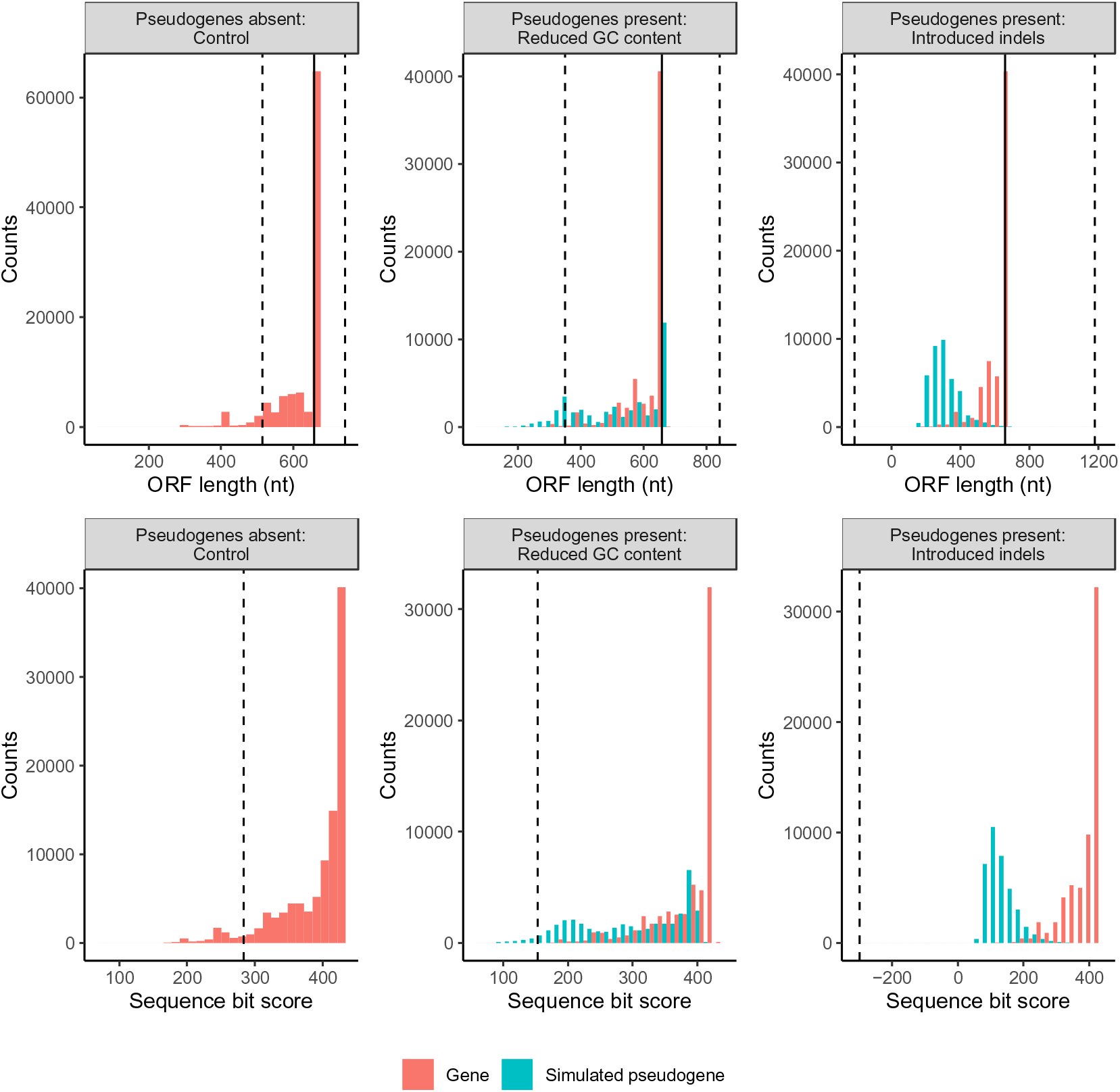
Doubling the proportion of mutated sequences greatly reduces the number of simulated pseudogenes removed. Each column shows the results from a particular simulation: a controlled community with pseudogenes absent, a community with pseudogenes that have a reduced GC content, and a community with pseudogenes where we have introduced indels. The top panel shows the length variation of sequences in the longest retained open reading frame. The solid vertical line indicates the length of a typical COI barcode at 658 bp. The two vertical dashed lines shows the boundaries for identifying ORFs with outlier lengths. The bottom panel shows the sequence bit score variation. The vertical dashed line shows the boundary for identifying sequences with small outlier scores.

**Fig S15.**
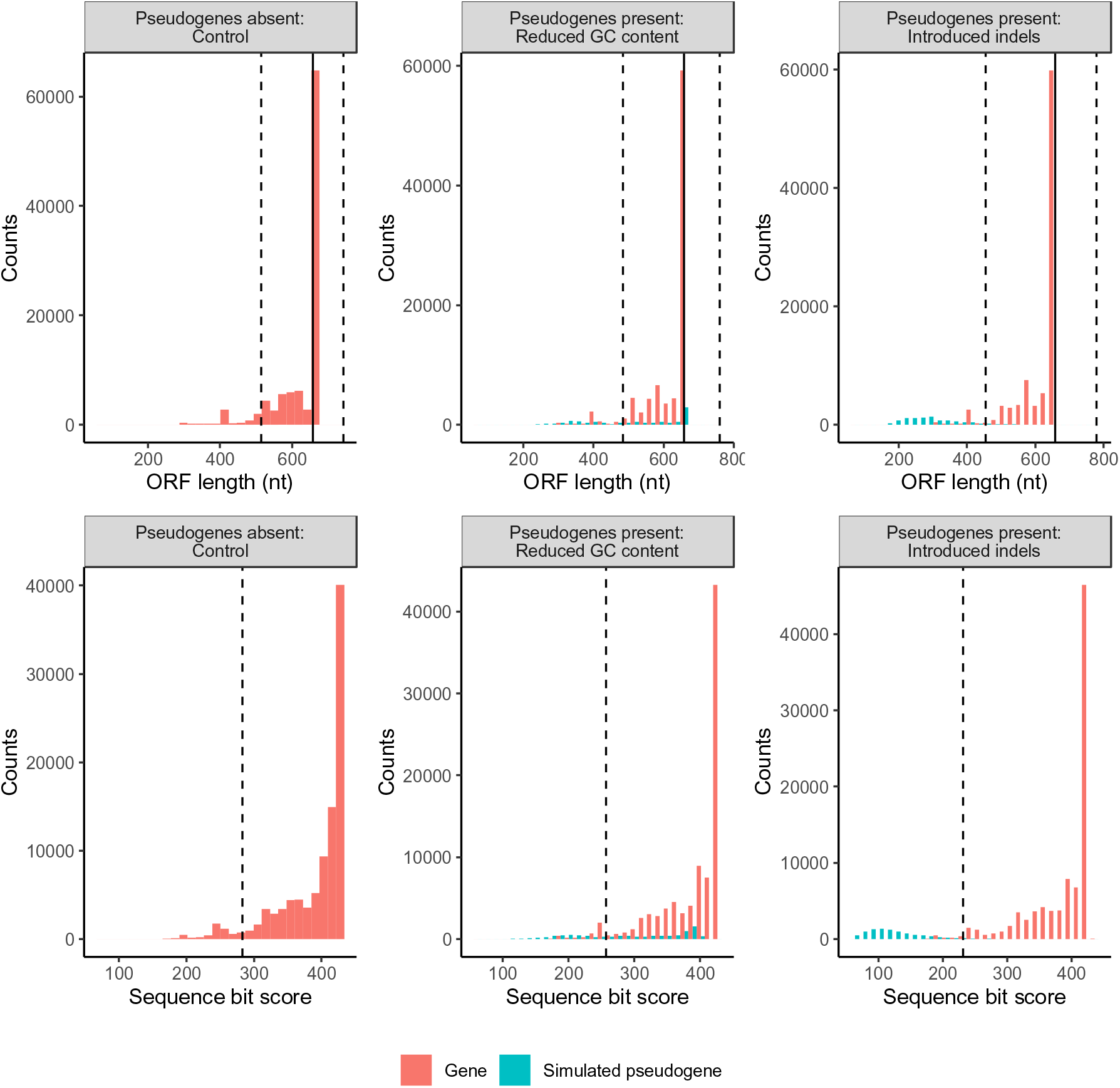
Halving the proportion of mutated sequences increases the number of simulated pseudogenes removed. Each column shows the results from a particular simulation: a controlled community with pseudogenes absent, a community with pseudogenes that have a reduced GC content, and a community with pseudogenes where we have introduced indels. The top panel shows the length variation of sequences in the longest retained open reading frame. The solid vertical line indicates the length of a typical COI barcode at 658 bp. The two vertical dashed lines shows the boundaries for identifying ORFs with outlier lengths. The bottom panel shows the sequence bit score variation. The vertical dashed line shows the boundaries for identifying sequences with short outliers scores.

